# DNA methylation restricts coordinated germline and neural fates in embryonic stem cell differentiation

**DOI:** 10.1101/2022.10.22.513040

**Authors:** Mathieu Schulz, Aurélie Teissandier, Elena de la Mata, Mélanie Armand, Julian Iranzo, Fatima El Marjou, Pierre Gestraud, Marius Walter, Sarah Kinston, Berthold Göttgens, Maxim V.C. Greenberg, Deborah Bourc’his

**Affiliations:** Institut Curie, PSL Research University, INSERM U934, CNRS UMR3215, Paris, France; Institut Curie, PSL Research University, INSERM U900, MINES ParisTech, Paris, France; Fred Hutchinson Cancer Center, Seattle, USA; Wellcome-MRC Cambridge Stem Cell Institute, University of Cambridge, Cambridge, UK; Université Paris Cité, CNRS, Institut Jacques Monod, F-75013 Paris, France

## Abstract

Somatic DNA methylation is established early during mammalian development, as embryonic cells transition from naive to primed pluripotency. This precedes the emergence of the three somatic germ layers, but also the segregation of the germline that undergoes genome-wide DNA demethylation after specification. While DNA methylation is essential for embryogenesis, the point at which it becomes critical during differentiation and whether all lineages equally depend on it is unclear. Using culture modeling of cellular transitions, we found that DNA methylation-free embryonic stem cells (ESCs) with a triple DNA methyltransferase knockout (TKO) normally progressed through the continuum of pluripotency states, but demonstrated skewed differentiation abilities towards neural versus other somatic lineages. More saliently, TKO ESCs were fully competent for establishing primordial germ cell-like cells (PGCLCs), even showing temporally extended and self-sustained capacity for the germline fate. By mapping chromatin states, we found that the neural and germline lineages are linked by a similar enhancer dynamics during priming, defined by common sets of methyl-sensitive transcription factors that fail to be decommissioned in absence of DNA methylation. We propose that DNA methylation controls the temporality of a coordinated neural-germline axis of preferred differentiation route during early development.

## INTRODUCTION

Cytosine DNA methylation is an epigenetic mark associated with transcriptional repression, providing critical control over development as illustrated by the embryonic lethality of DNA methyltransferase (*Dnmt*) mutant mice ^1,2^. DNA methylation is highly dynamic in early embryonic cells, coinciding with functional changes in potency ^3^. Pluripotent stem cells of the blastocyst have globally low genomic methylation levels, associated with unbiased ability to form all embryonic lineages. Somatic DNA methylation patterns are established during epiblast formation and prior to gastrulation, upon transitioning to the primed pluripotent state ^4,5^. During naive-to-primed pluripotency transition, an intermediate formative state allows for an important fate bifurcation towards primordial germ cells (PGCs) ^6–8^. PGC specification is followed by global erasure of somatic DNA methylation, allowing for the subsequent establishment of germline-specific patterns ^9^.

Maintenance and differentiation of embryonic stem cells (ESCs) in culture can provide key insights into early development and its associated DNA methylation dynamics. For example, DNA methylation-free *Dnmt1*, *Dnmt3A* and *Dnmt3B* triple-knockout (*Dnmt*-TKO) ESCs can be maintained in naive-prone conditions, but they have an impaired ability for somatic differentiation ^10–12^. There are several manners through which DNA methylation could directly influence early cell state transitions. To wit, during epiblast differentiation, some naive pluripotent genes acquire DNA methylation at their promoters and/or enhancers ^13,14^, which may indicate a role in pluripotency dissolution and priming for differentiation ^12^. Moreover, lineage-specific enhancers display heterogeneous DNA methylation levels in populations of naive and primed ESCs in culture ^15–17^, which may reflect requisite intercellular heterogeneity as cells prepare to engage towards specific lineages. Mechanistically, DNA methylation anti-correlates with chromatin features that reflect enhancer activity, such as lower nucleosome occupancy and histone 3 lysine 27 acetylation (H3K27ac) ^18^, possibly by restraining the binding of DNA methylation-sensitive transcription factors (TFs) ^19,20^.

Here, we examined the function of DNA methylation in critical cell fate bifurcations, namely pluripotency state transition and somatic versus germline commitment. Using *Dnmt*-TKO ESCs, we found that DNA methylation is dispensable for acquiring transcriptional and functional features associated with priming, *in vitro* and *in vivo*. Regarding differentiation potential, DNA methylation-free ESCs exhibited predilection for entering the neural fate versus other somatic lineages. Perhaps more strikingly, lack of DNA methylation provided extended temporal opportunity and autonomous competence for germline induction. By mapping chromatin accessibility and activity, we found that the propensity towards both neural and germline lineages was linked to a failure to decommission putative neural and germline enhancers that were co-regulated by similar chromatin dynamics and DNA methylation-sensitive transcription factors. We propose that DNA methylation is required at a subset of DNA methylation-sensitive enhancers during priming to limit an early coordinated neutral-germline fate as preferred route of differentiation.

## RESULTS

DNA methylation is dispensable for naive-to-primed pluripotency transition DNA methylation is dispensable for naive pluripotent cells in culture ^10^. Whether it plays a role upon transitioning from naive to primed pluripotency has not been addressed, although this functional switch coincides with genome-wide acquisition of DNA methylation ^3,14^. To probe the role of DNA methylation, we engineered a new *Dnmt-*TKO line from low passage ESCs (E14 background) (Extended Data Fig. 1), to mitigate effects due to potential genetic and epigenetic drift upon prolonged time culture. To recapitulate the naive-to-primed transition, we performed *in vitro* Epiblast-like cell (EpiLC) differentiation, which includes a transient phase of formative pluripotency at 40-48h ^5,6,21^ (Fig. 1a). While both wild-type (WT) and TKO ESCs had low genomic methylation content in 2i/LIF+Vitamin C medium ^22^ at day 0 (D0), they diverged upon EpiLC differentiation: WT EpiLCs reached over 71% of CpG methylation at D4, while TKO EpiLCs stayed DNA methylation-free (Fig. 1b). However, TKO EpiLCs displayed similar morphology, growth and apoptosis rate than WT EpiLCs (Fig. 1c and Extended Data Fig. 2a,b).

**Fig. 1.**
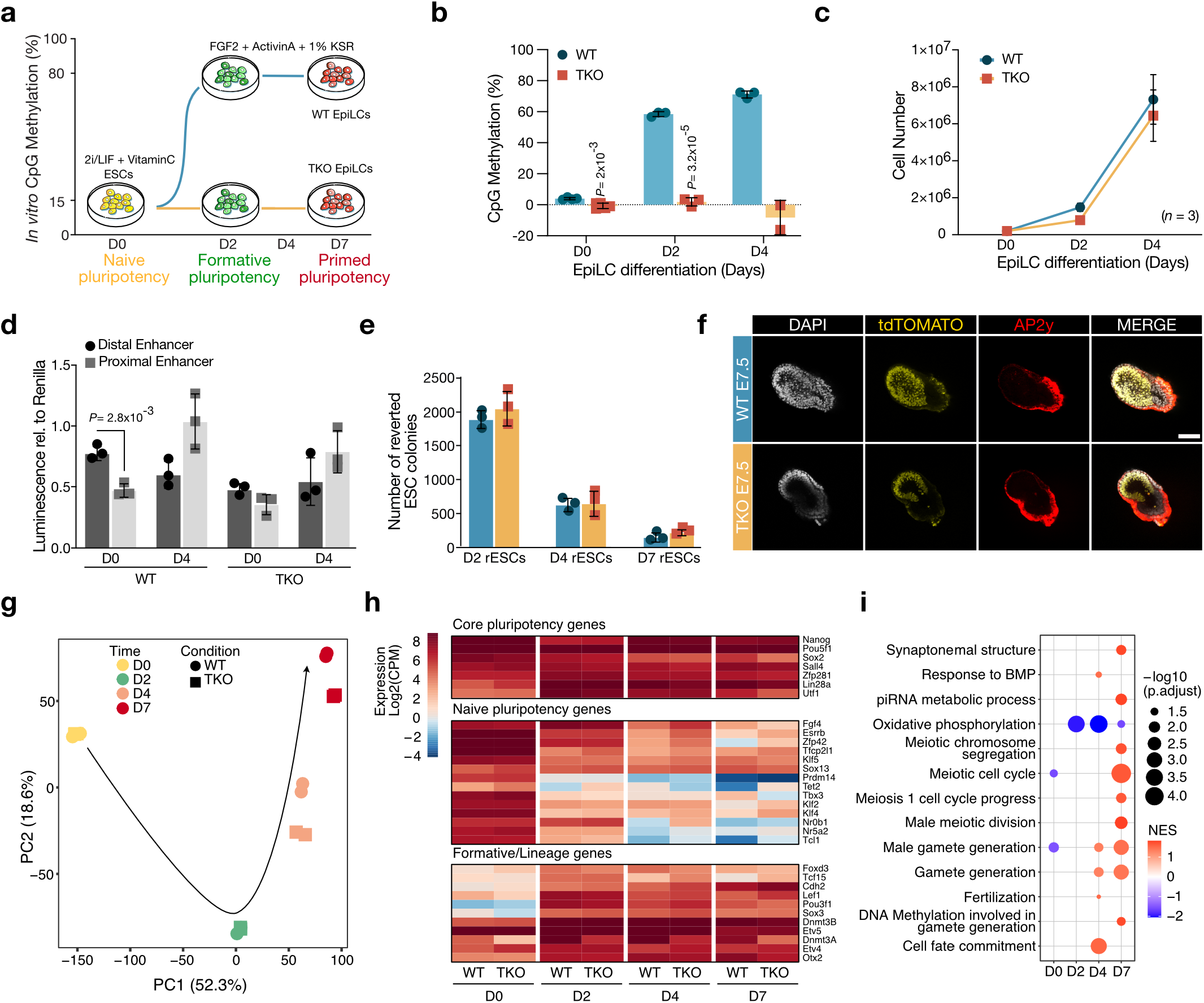
TKO ESCs are competent for naive-to-primed pluripotency transition. **a,** Schematic representation of Epiblast-like cell (EpiLC) differentiation. Axes represent genomic methylation levels and key pluripotency states across time. **b,** Barplot representing CpG methylation percentage during WT and TKO EpiLC differentiation as measured by LUMA. Data are mean ± SD from biological replicates (*n=* 3 or 2) (two-tailed unpaired student t-test). **c,** Growth curve during WT (blue) and TKO (orange) EpiLC differentiation. Data are from biological triplicates with mean ± SD. **d,** Barplot representing Luciferase activity from the distal (dark grey) and proximal (light grey) *Oct4* enhancers in WT and TKO D0 ESCs and D4 EpiLCs. Data are mean ± SD from biological triplicates (two-tailed unpaired student t-test). **e,** Barplot displaying number of reverted ESC colonies upon 2i+VitC medium replacement at D2, D4 and D7 of EpiLC differentiation. Data are from biological triplicates with mean ± SD. **f,** Confocal images of E7.5 chimeric embryos obtained by morula aggregation with WT or TKO H2B::tdTOMATO-transgenic ESCs. Embryos were stained for DAPI (gray) and AP2γ (red, extra-embryonic marker). Both WT and TKO H2B::tdTOMATO cells showed strong contribution to the epiblast, but not extra-embryonic tissues. Scale: 100µm. **g,** Principal component analysis (PCA) from bulk RNA-seq of WT (circles) and TKO (squares) EpiLCs from D0 to D7 of differentiation in biological duplicates. Axes indicate two main components for variability, the black arrow represents hypothetical trajectory. **h,** Heatmaps of pluripotency marker expression showing normalized Log2 CPM counts during EpiLC differentiation (average between biological duplicates). **i,** Gene set enrichment analysis (GSEA) showing normalized enrichment score (NES) and -log10(p.adj) for germline-associated ontologies in TKO over WT EpiLC differentiation (pvalue cutoff= 0.05).

To capture the identity of TKO EpiLCs, we assessed their functional features. First, we performed a dual luciferase assay that measures the *Oct4* enhancer switch that occurs upon priming ^16^. We observed activation of the proximal enhancer in both WT and TKO EpiLCs at D4, compared to D0 naive ESCs (Fig. 1d), indicating proper priming. Next, we tested the propensity of TKO EpiLCs to revert towards a naive state upon changing for 2i/LIF+Vitamin C medium. The absence of DNA methylation did not confer higher reversion: both WT and TKO yielded similar, and progressively decreasing numbers of alkaline phosphatase-positive reverted ESC (rESC) colonies from D2 to D7 EpiLCs (Fig. 1e). Finally, we tested whether TKO ESCs could contribute to the post-implantation epiblast *in vivo*. For this, we aggregated WT or TKO ESCs expressing a constitutive H2B::tdTOMATO reporter, with host morulae (Extended Data Fig. 2c-f). Upon implantation in foster females, TKO H2B::tdTOMATO ESCs showed contribution to the epiblast of E7.5 chimeric embryos, similarly to WT (Fig. 1f and Extended Data Fig. 2g), supporting our *in vitro* findings that DNA methylation is dispensable for acquiring primed epiblast features. However, TKO chimerae were delayed and smaller than WT chimerae, indicating that DNA methylation may become necessary for epiblast development after priming. While a previous study showed trophectoderm differentiation of double *Dnmt3A; Dnmt3B* KO ESCs ^23^ after blastocyst injection, we did not observe extra-embryonic contribution of the triple *Dnmt* KO ESCs. Accordingly, the trophectoderm marker *Ascl2* was not up-regulated during TKO EpiLC differentiation *in vitro* (Extended Data Fig. 2h).

Bulk RNA sequencing (RNA-seq) followed by principal component analysis (PCA) further highlighted the transcriptional similarity of D0 ESCs and D2 EpiLCs between WT and TKO backgrounds (Fig. 1g and Supplementary Table 1). In particular, naive markers were properly repressed and primed markers induced in D2 TKO EpiLCs (Fig. 1h). However, stronger divergence was observed at later timepoints: TKO EpiLCs demonstrated 1,610 and 2,687 mis-regulated genes at D4 and D7, respectively, the majority being up-regulated with an overrepresentation of genes expressed in the germline (Fig. 1i, Extended Data Fig. 3a,b and Supplementary Table 2), which are known to be regulated by promoter DNA methylation ^24^. In particular, we found that among the 581 genes that gained promoter DNA methylation (>75%) in WT EpiLCs and were up-regulated in D7 TKO EpiLCs, 103 were germline genes (Extended Data Fig. 3c,d). Various families of transposable elements (TEs) were also over-expressed in TKO EpiLCs (Extended Data Fig. 3e), with Intracisternal A Particle (IAP) elements as early as D2 of differentiation, and Long Interspersed elements (LINE-1) at D4.

In sum, we found that despite increasing during naive-to-primed pluripotency transition, DNA methylation appears dispensable for the other hallmarks of priming, *in vitro* and *in vivo*. To exclude potential clonal effects, we confirmed that priming is decoupled from DNA methylation with an independent TKO E14 clone (Extended Data Fig. 3f,g) ^25^. This raises the question as to when DNA methylation becomes essential during the ensuing steps of somatic development.

### DNA methylation restricts neural routing during somatic differentiation

Previous studies yielded contrasting results regarding the viability and ability of TKO ESCs to properly activate somatic transcriptional programs during *in vitro* differentiation ^11,12^, despite using the same originally derived TKO ESC line ^10^. To shed light onto the role on DNA methylation in the acquisition of somatic identities, we first performed undirected differentiation towards embryoid bodies (EBs) with our newly derived TKO ESC line. While WT EBs continuously grew during the eight days of differentiation, TKO EBs stopped expanding after D3 (Extended Data Fig. 4a,b), independently of initial cell density (Extended Data Fig. 4c,d). DNA methylation appears therefore critical for multi-lineage somatic differentiation in EBs.

To probe the influence of DNA methylation in adopting a specific somatic fate, we then performed directed neural progenitor cell (NPC) differentiation ^26^. WT cells demonstrated strong growth rate during the 10 days of NPC differentiation, adopting neural-like morphological features such as axon-like protrusions and neural rosettes (Fig. 2a,b). TKO cells survived NPC differentiation, although showing a lower growth rate associated with lower viability from D8 and onwards, and formed protrusions but not more advanced structures (Fig. 2a-d). PCA of bulk RNA-seq performed across NPC differentiation revealed dominant ordering of cells by time (PC1, 58.1%), whereas the absence of DNA methylation only accounted for 12.5% (PC2) of total variance despite the up-regulation of germline genes and TEs (notably IAP and LINE-1) in TKO NPCs (Fig. 2e and Extended Data Fig. 4e,f). While neurogenic gene ontology (GO) terms were first up-regulated in D2 TKO compared to WT NPCs, most of these terms became under-represented at D8, reflecting secondary neurogenic gene down-regulation (Fig. 2f,g and Supplementary Table 2). Simultaneously, the pluripotency network became abnormally reactivated in late TKO NPCs at D8, which was associated with a “U-turn” in the PCA plot (Fig. 2h and Supplementary Table 2). DNA methylation is therefore not required for inducing a neural identity, but it becomes necessary in the long-term to stabilize this identity and/or to prevent spontaneous dedifferentiation.

**Fig. 2.**
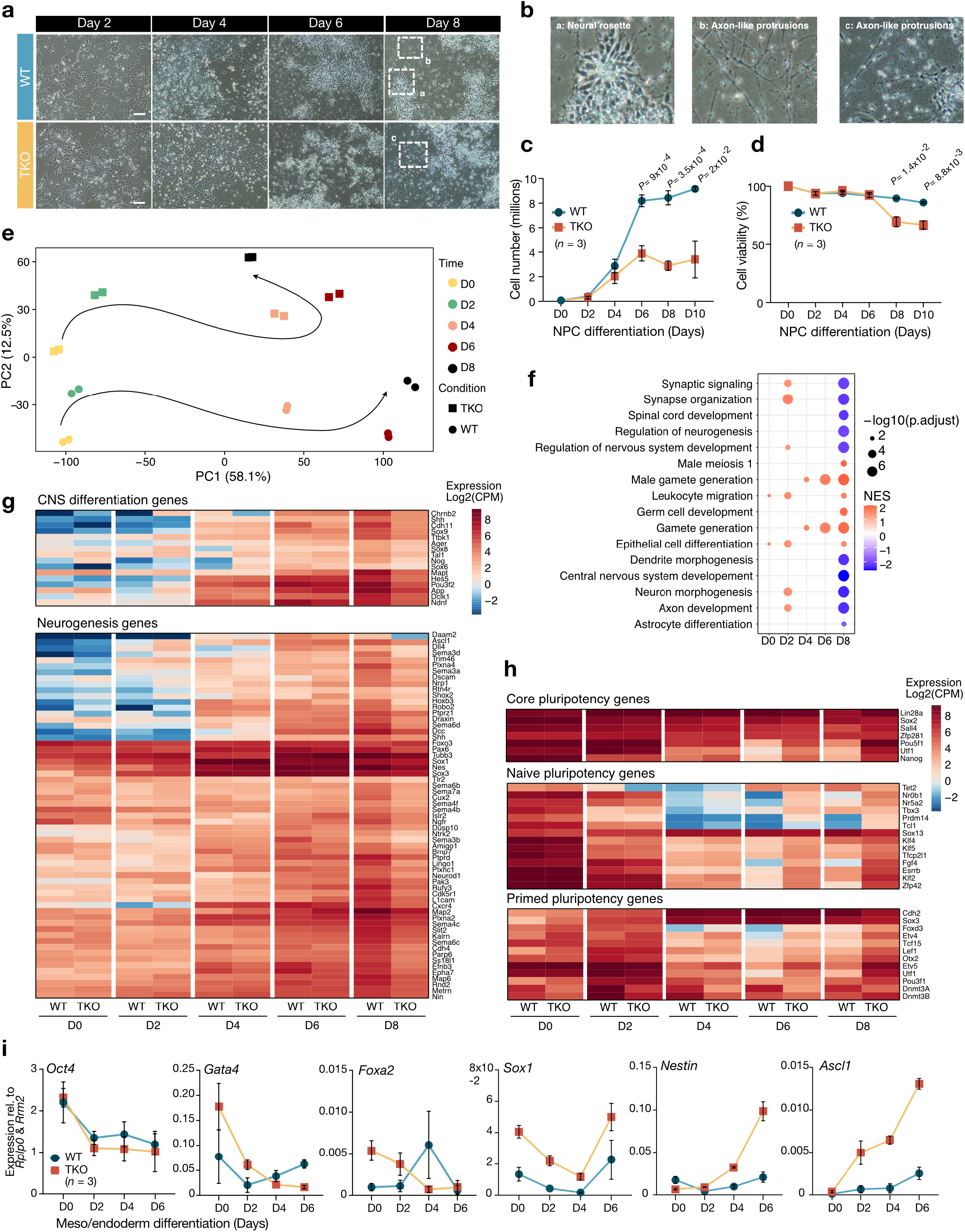
TKO ESCs can induce a neural fate but fail to consolidate long-term repression of pluripotency. **a, b,** Brightfield images of (**a**) neural progenitor cells (NPCs) generated from WT and TKO ESCs cultured in N2B27 for 8 days. Scale: 200µm. (**b**) White dotted squares are a close-up view of: a. Neural rosettes, b and c. Axon-like protrusions. **c, d,** (**c**) Growth and (**d**) Trypan blue-based viability curves upon WT and TKO NPC differentiation. Data are mean ± SD in biological triplicates (two-tailed unpaired student t-test). **e,** PCA from bulk RNA-seq during WT (circles) and TKO (squares) NPC differentiation in biological duplicates. Axes indicate two main components for variability, the black arrow represents hypothetical trajectory. **f,** Gene set enrichment analysis (GSEA) showing normalized enrichment score (NES) and -log10(p.adj) for neural and germline associated-ontologies in TKO over WT NPC differentiation (pvalue cutoff=0.05). **g, h,** Heatmaps of (**g**) Neurogenesis and (**h**) Pluripotency markers during NPC differentiation. Expression level was obtained for each condition from the average between biological duplicates and represented as normalized Log2 CPM counts. **i,** Expression of pluripotency (*Oct4*), meso/endoderm (*Gata4*, *Foxa2*) and neural (*Sox1*, *Nestin*, *Ascl1*) markers measured by RT-qPCR in WT and TKO during mesoderm differentiation. Data shown are mean ± SD from biological triplicates. ΔCT values were normalized to *Rplp0* and *Rrm2* (two-tailed unpaired student t-test).

We tested another somatic differentiation route *in vitro*, towards the meso/endoderm lineage ^23^. TKO cells survived the protocol (Extended Data Fig. 4g) and repressed the pluripotency marker *Oct4* (Fig. 2i). However, they failed to properly activate the meso/endoderm markers *Gata4* or *Foxa2*, and rather up-regulated neural genes, such as *Sox1*, *Nestin* and *Ascl1* (Fig. 2i). Therefore, upon somatic *in vitro* differentiation, the lack of DNA methylation may favor the neural lineage over other somatic lineages.

### DNA methylation restricts the temporal window of germline specification

We then addressed the role of DNA methylation in the bifurcation between germline and somatic fates when cells transit throughout the formative phase. DNA methylation-free TKO cells could harbor higher competence for germline induction, as primordial germ cells (PGCs) naturally acquire an hypomethylated genome following their specification and re-express genes typical of naive pluripotency ^9,27^.

We used PGC-like cell (PGCLC) differentiation to generate *in vitro* germ cell precursors from EpiLCs at 40h of differentiation (Fig. 3a), which is considered as the optimal timing for PGCLC specification ^28,29^. TKO EpiLCs were able to form and grow as aggregates under germline-driving conditions (Extended Data Fig. 5a). Specified PGCLCs were sorted using the SSEA1 and CD61 surface markers (Fig. 3b and Extended Data Fig. 5b), and RT-qPCR validated similar induction of early PGCLC markers in WT and TKO double-positive compared to double-negative cells (Fig. 3c and Extended Data Fig. 5c). PGCLC specification rates were globally similar between WT and TKO population of cells (15.3% and 17.3%, respectively) (Fig. 3d), indicating that TKO cells can undergo germline specification *in vitro*. We then investigated whether DNA methylation could condition the specification timing of the germline during epiblast progression. As previously reported ^30^, both WT and TKO naive ESCs were not adequately competent to give rise to PGCLCs directly (Fig. 3d). During WT EpiLC differentiation, germline competency was progressively gained until 40h (15.3% of specified PGCLCs), before dropping at 48h and 96h (3.1% of specified WT PGCLCs at 96h), as late EpiLCs become fully primed for somatic lineages. In contrast, TKO EpiLCs maintained germline competence beyond 40h. Notably, 96h TKO EpiLCs were four times more efficient for PGCLC specification than time-matched WT EpiLCs (12.4% of specified TKO PGCLCs) (Fig. 3d and Extended Data Fig. 5d), which we confirmed in the independent E14 TKO clone (Extended Data Fig. 5e). This suggests that the lack of DNA methylation provides a protracted window for adopting the germline fate. Moreover, when we induced PGCLC differentiation without adding pro-germline cytokines, like BMPs and others, we found that TKO EpiLCs exhibited autonomous ability to give rise to the germline, while WT EpiLCs were poorly competent (Extended Data Fig. 5f).

**Fig. 3.**
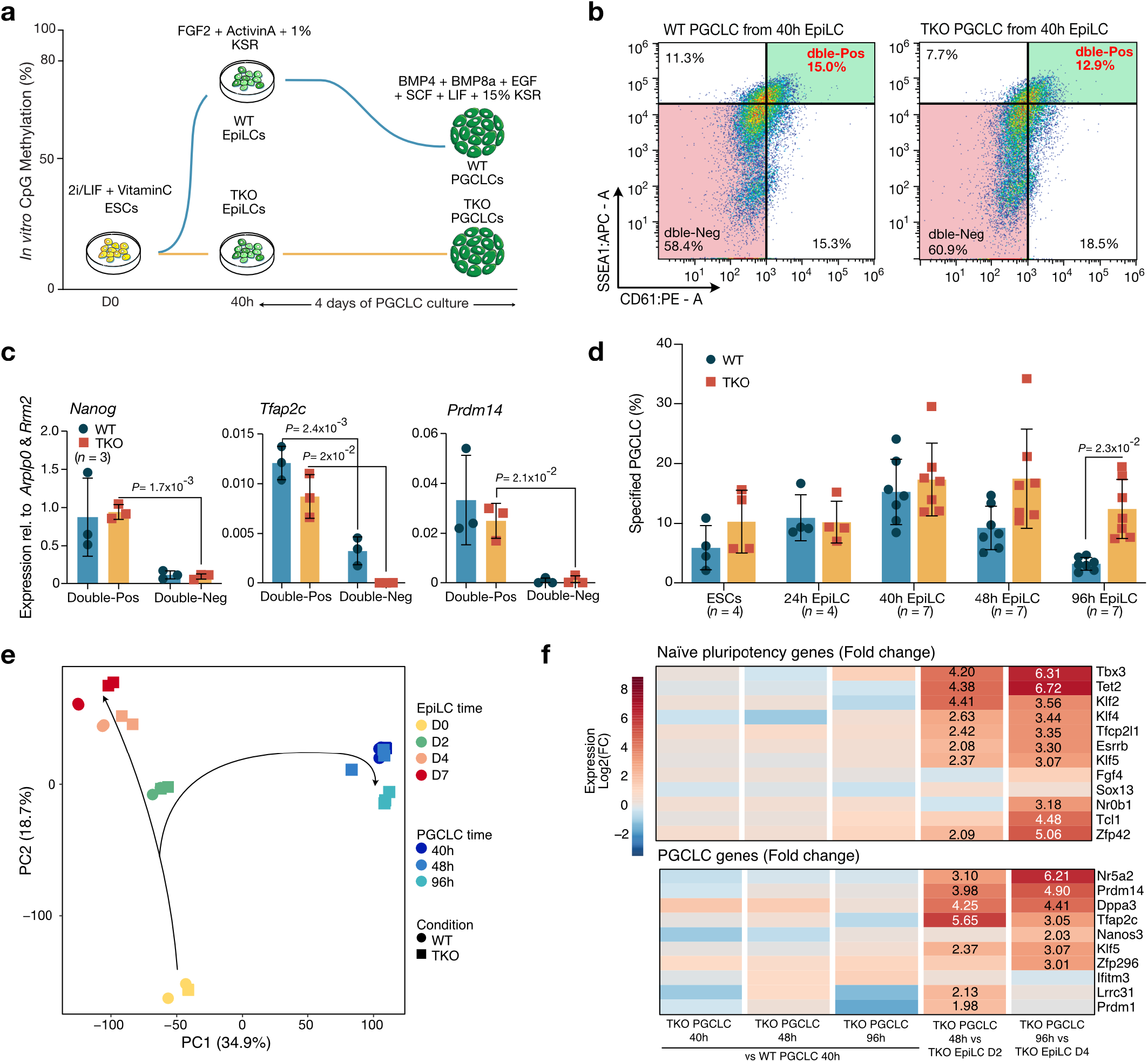
TKO EpiLCs demonstrate longer opportunities for germline contribution *in vitro.* **a,** Schematic representation of PGC-like cell (PGCLC) differentiation, with axes representing genomic methylation levels and time. **b,** Representative FACS plot of double SSEA1-/ CD61-Positive WT and TKO PGCLCs generated from 40h EpiLCs. Viable cell percentages are indicated in each quarter. **c,** Expression of PGCLC markers measured by RT-qPCR in WT (blue) and TKO (orange) PGCLCs. Data shown are mean ± SD from biological triplicates as ΔCT values normalized to *Arplp0* and *Rrm2* (two-tailed unpaired student t-test). **d,** Barplot showing time-window percentage of specified WT and TKO PGCLCs generated from ESCs and EpiLCs at different times. Data shown are mean ± SD from biological replicates (*n*=4 or 7) (two-tailed unpaired student t-test). **e,** PCA from bulk RNA-seq of WT (circles) and TKO (squares) EpiLC and PGCLC differentiation in biological duplicates. Axes indicate variance, black arrows represent hypothetical trajectory. **f,** Heatmaps of naive pluripotency and PGCLC markers showing normalized Log2 CPM counts Fold change of TKO over WT PGCLCs, and TKO PGCLCs over TKO EpiLCs. Numbers are indicated when log2FC>2. Expression was obtained for each condition from the average between biological triplicates.

Using bulk RNA-seq, we observed that WT and TKO PGCLCs clustered closely together in the PCA, highlighting high transcriptomic similarity, independently of the timing at which TKO PGCLCs were induced during EpiLC differentiation (Fig. 3e and Supplementary Table 1). Focusing on naive pluripotency and PGCLC markers, late induced TKO PGCLCs demonstrated similar activation compared to canonical WT PGCLCs generated from 40h EpiLCs, and expected reactivation compared to their corresponding D2 and D4 EpiLC precursors (Fig. 3f). This provides strong evidence that late TKO EpiLCs generate *bona fide* PGCLCs.

Previous studies have proposed that retention of naive pluripotency during differentiation could prime cells towards germline specification ^31,32^. To check whether TKO EpiLCs could be delayed in exiting naive pluripotency, we performed single cell RNA-seq (SMART-seq2) at critical time-points (D0, D1, D2 and D4). We did not detect specific TKO EpiLC subpopulations compared to WT (Extended Data Fig. 6a,b). Clustering analysis revealed five clusters—termed “Naive”, “Exit Naivety”, “Transitioning”, “Formative” and “Primed” based on their gene expression profile— that mostly overlaid with EpiLC differentiation days (Extended Data Fig. 6c,d and Supplementary Table 3). Noticeably, the “Primed” cluster demonstrated partition between WT and TKO cells, which we attributed to the germline gene up-regulation in D4 TKO EpiLCs (Extended Data Fig. 6b,e). When focusing on key markers of early PGC induction *in vivo* or *in vitro*, we did not observe clusters with higher expression of these genes in TKO cells (Extended Data Fig. 6f). Finally, pseudotime analysis did not reveal asynchronous or diverging trajectory upon TKO EpiLC differentiation (Extended Data Fig. 6g).

Overall, these results demonstrate that the absence of DNA methylation extends the window of germline specification during EpiLC differentiation and provides self-sustained ability for committing to a PGC fate, but this was not due to persistence of the naive pluripotency gene program.

### DNA methylation counteracts chromatin accessibility during EpiLC differentiation, without directly regulating gene promoters

Concomitantly with genome-wide acquisition of DNA methylation, the chromatin landscape is globally remodeled during naive-to-primed pluripotency transition. In particular, it was shown that chromatin accessibility and H3K27ac levels are profoundly remodeled at enhancers and promoters of pluripotency and developmental genes ^18,21,33^. To investigate whether failure to gain DNA methylation could impact the priming-associated chromatin program, we assessed chromatin accessibility by Assay for Transposase-Accessible Chromatin (ATAC-seq) and H3K27ac distribution by Cleavage Under Targets and Release Using Nuclease (CUT&RUN) in D0 ESCs and D2-D4 EpiLCs, when TKO cells display extended abilities for PGCLC differentiation (Supplementary Table 1).

At the global level, D2 and D4 TKO EpiLCs demonstrated abnormal retention of sharp ATAC and H3K27ac-enriched regions, while these tended to decline across WT EpiLC differentiation (Fig. 4a,b and Supplementary Tables 4-5). More precisely, using Differential Enrichment (DE) analysis, 11,079 and 14,525 regions demonstrated significant ATAC or H3K27ac increase, respectively, in D4 TKO compared to WT EpiLCs (Fig. 4c). Among these, 47% of ATAC-increased regions and 26% of H3K27ac-increased regions mapped to promoters (<5kb from gene transcription start site, TSS) (Fig. 4d). However, only few differentially expressed genes were linked to promoter-related ATAC or H3K27ac changes in D4 TKO EpiLCs (486 and 187 mis-regulated genes, respectively, see Supplementary Tables 4 and 5) (Extended Data Fig. 7a,b). Moreover, these promoters showed scarce DNA methylation gain during WT EpiLC differentiation as assessed from Whole Genome

**Fig. 4.**
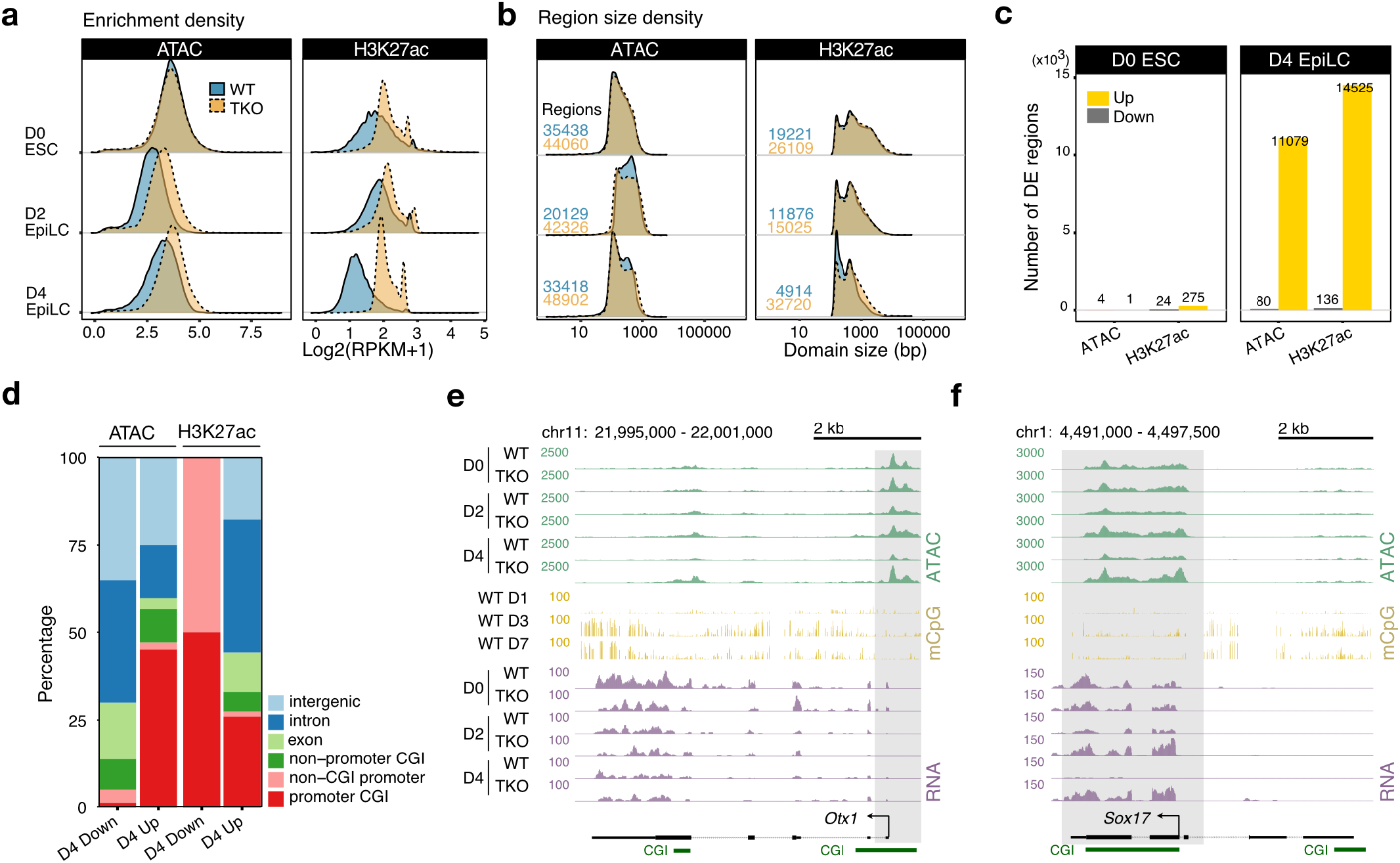
TKO EpiLCs display a globally relaxed chromatin. **a, b,** Ridge plots displaying global (**a**) Enrichment density in Log2(RPKM+1) and (**b**) Region size density (in bp) plots, for ATAC- and H3K27ac-marked regions in WT (blue) and TKO (orange) EpiLCs at D0, D2 and D4 of differentiation. **c,** Differential Enrichment (DE) analysis of ATAC and H3K27ac enrichment in TKO over WT cells. Number of gained regions (“up”, yellow) and lost regions (“down”, grey”) are shown at D0 and D4 of EpiLC differentiation (FDR<0.05). **d,** Genomic annotation analysis of DE regions in D4 EpiLCs. **e, f,** UCSC tracks showing increased ATAC (green) signal in D4 TKO EpiLCs (**e**) with no impact on gene expression (purple, RNA-seq) or (**f**) with an impact on gene expression. In both situations, (**e**) *Otx1* and (**f**) *Sox17* show an hypomethylated (yellow, WGBS) CGI promoter in WT EpiLCs.

Bisulfite Sequencing (WGBS) data ^34^ (<25% of CpG methylation at D7) (Fig. 4e,f and Extended Data Fig. 7c,d), indicating independent effects from DNA methylation-based promoter silencing. Overall, failure to remethylate the genome induced retention of a globally relaxed chromatin during priming—both in terms of accessibility and histone marks—but without major and direct impact on gene promoter regulation.

### DNA methylation contributes to NPC and PGCLC enhancer regulation during priming

To understand why TKO cells showed preferred ability to enter the neural lineage and extended competence for germline differentiation, we next specifically focused on NPC and PGCLC enhancers. We defined putative NPC (*n*=168) or PGCLC enhancers (*n*=350) as regions with overlapped H3K27ac and ATAC signals at any time point of WT EpiLC differentiation, located in a 50kb window of known NPC- or PGCLC-specific genes (genes were declared NPC- or PGCLC-specific if they demonstrated FDR<0.05 and Log2CPM>1 in our RNA-seq datasets, see Supplementary Table 2). Co-occurrence of these features is considered as a hallmark of enhancer activity ^35,36^. These putative NPC or PGCLC enhancers abnormally retained H3K27ac and/or ATAC signals in D2 and D4 TKO EpiLCs over WT (Fig. 5a-c and Supplementary Table 6). Importantly, enhancers associated with endoderm and mesoderm development ^37^ were not enriched in ATAC and H3K27ac in TKO over WT EpiLCs (Fig. 5d), suggesting lineage-specific retention of active chromatin signatures at NPC and PGCLC enhancers.

**Fig. 5.**
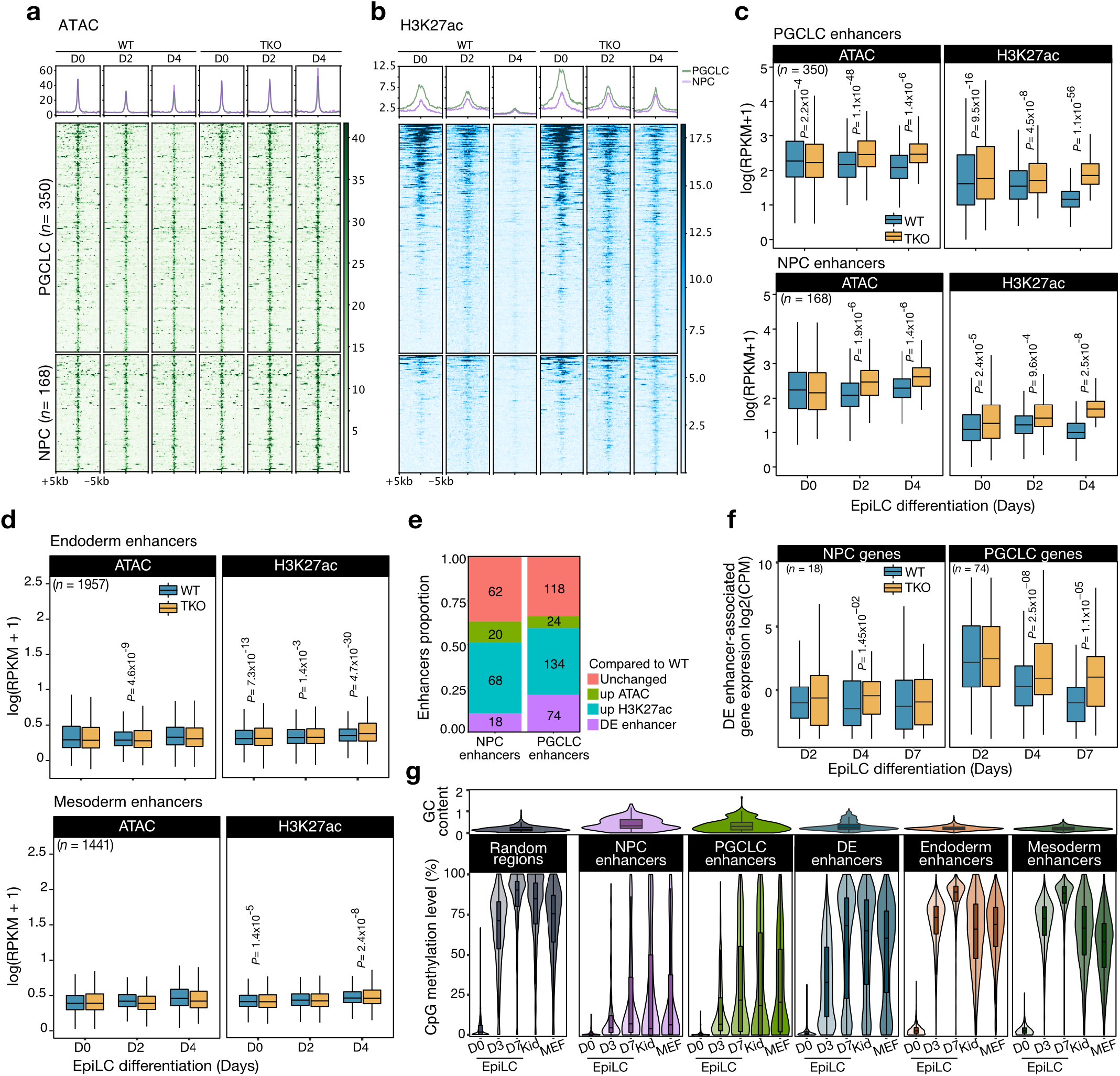
TKO EpiLCs retain active chromatin marks specifically at NPC and PGCLC putative enhancers. **a, b,** Heatmap and metaplot for (**a**) ATAC and (**b**) H3K27ac for PGCLC (green, *n*=350) and NPC (purple, *n*=168) putative enhancers in WT and TKO differentiating EpiLCs. Enrichment in RPKM for 10kb windows around the centered regions are represented. **c,** Boxplot showing enrichment in Log(RPKM+1) of ATAC and H3K27ac at PGCLC and NPC putative enhancers, for WT (blue) and TKO (orange) differentiating EpiLCs. Data shown are the median with upper and lower hinges corresponding to 75 and 25% quantile (two-tailed unpaired student t-test). **d,** same as **c**) but at previously defined endoderm (*n*=1957) and mesoderm enhancers (*n*=1441) ^37^. **e,** Barplot showing the proportion and absolute number of putative enhancers in D4 TKO EpiLCs gaining ATAC or H3K27ac, or being differentially enriched (DE) in both marks when compared to WT. **f,** Boxplot showing expression in Log2(CPM) of genes associated with NPC (*n*= 18) or PGCLC (*n*= 74) DE enhancers, for WT and TKO cells during EpiLC differentiation. Data shown are the median with upper and lower hinges corresponding to 75 and 25% quantile (two-tailed paired student t-test). **g,** DNA methylation analysis at enhancers. Top panel: global GC content as a ratio of observed/expected GC nucleotides. Lower panel: dynamics of methylated CpGs during WT EpiLC differentiation (D0, D3, D7), in kidney (Kid) and in Mouse Embryonic Fibroblasts (MEFs) at previously defined enhancers compared to random regions, using public WGBS datasets ^34,41,42^. Data shown are the median with upper and lower hinges corresponding to 75 and 25% quantile.

Then, we considered all genomic regions that significantly gained both ATAC and H3K27ac signals in D4 TKO EpiLCs, which we termed differentially enriched enhancers (DE enhancers). We focused on D4 EpiLCs, since this time-point coincides with the strongest chromatin effects and highest bias for adopting a germline fate in TKO versus WT cells. We identified 1,339 putative DE enhancers scattered in the genome of D4 TKO EpiLCs, among which 18 and 74 coincided with previously defined NPC and PGCLC enhancers, respectively (Fig. 5e). Gene up-regulation in TKO EpiLCs was observed in the vicinity of NPC- and PGCLC-associated DE enhancers from D4 and later on, among which were critical genes involved in priming and PGCLC specification, such as *Zic3*, *Etv5*, *Dppa2*, *Dppa4* and *Zfp42* ^38–40^ (Fig. 5f and Supplementary Table 6). The other, non-NPC or PGCLC-associated DE enhancers, did not show significant up-regulation of nearby genes in TKO EpiLCs, NPCs or PGCLCs (Extended Data Fig. 8a), suggesting a potential pre-activation state.

Interestingly, NPC and PGCLC enhancers demarcated from random genomic regions by their higher GC content, a feature that was shared—although to a less extent—by DE enhancers. Using WGBS datasets ^34,41,42^, we found that NPC, PGCLC and DE enhancers acquired a wide range of DNA methylation levels during WT EpiLC differentiation, which was conserved in differentiated tissues (mouse embryonic fibroblasts MEFs, and kidney), compared to an homogeneously high gain of methylation genome-wide (Fig. 5g). Again, meso/endoderm enhancers behaved distinctively from NPC-PGCLC enhancers, with relatively low GC content and fast acquisition of high DNA methylation levels during EpiLC differentiation, similarly to the average genome (Fig. 5g), as previously observed in the post-implantation epiblast *in vivo* ^37^. Individual examination revealed that a subset of NPC, PGCLC and DE enhancers actually reached high DNA methylation levels during priming (18.3%, 30.8% and 64.7% of NPC, PGCLC and DE enhancers, respectively, in D7 EpiLCs) (Extended Data Fig. 8b and Supplementary Table 6), and these were related to critical genes involved in priming and/or neural and germline fates (*Ifitm3*, *Zic3*, *Etv5*, *Dppa2*, *Dppa4*, *Tet2*, *Tubb3* or *Sema*). DNA methylation-based regulation could therefore participate in decommissioning key NPC, PGCLC and DE enhancers during EpiLC differentiation.

In sum, PGCLC and NPC enhancers specifically retained H3K27ac and ATAC marks in differentiating TKO EpiLCs, while a subset of those enhancers normally gained high methylation levels in WT EpiLCs. Improper enhancer decommissioning was associated with up-regulation of a minimal number of NPC and PGCLC genes in TKO EpiLCs, but this might additionally poise these cells towards adopting neural or germline fates later on, when exposed to appropriate signals.

### DNA methylation reduces the binding of sensitive TFs at NPC and PGCLC enhancers

To identify which transcription factors (TFs) may drive the retention of NPC-PGCLC enhancer activity in TKO EpiLCs, we performed binding motif analysis using HOMER^43^. This revealed that PGCLC and NPC enhancers shared common TF binding motifs (Fig. 6a). Similar binding motifs were also found when considering all DE enhancers detected in TKO EpiLCs at D4, suggesting that PGCLC-NPC-DE enhancers might be co-regulated by the same TF sets (Fig. 6a). Moreover, identified TF binding motifs contained CpG dinucleotides, which are the substrates of DNA methylation (Fig. 6b). Accordingly, we noticed that many of these motifs related to factors that are regulated by or sensitive to DNA methylation, such as SP, KLF, CEBP, CTCF, ETS and BACH factors ^19,44–46^. In addition, we found motifs associated with TFs that regulate pluripotency transition, such as the pluripotency triad (OCT4, SOX2, NANOG), KLF, ZIC, ETS factors and ZFP281 ^39,47,48^. In comparison, meso/endoderm enhancers were strikingly regulated by distinct TFs sets, sharing very little overlap with NPC-PGCLC enhancers (Fig. 6c).

**Fig. 6.**
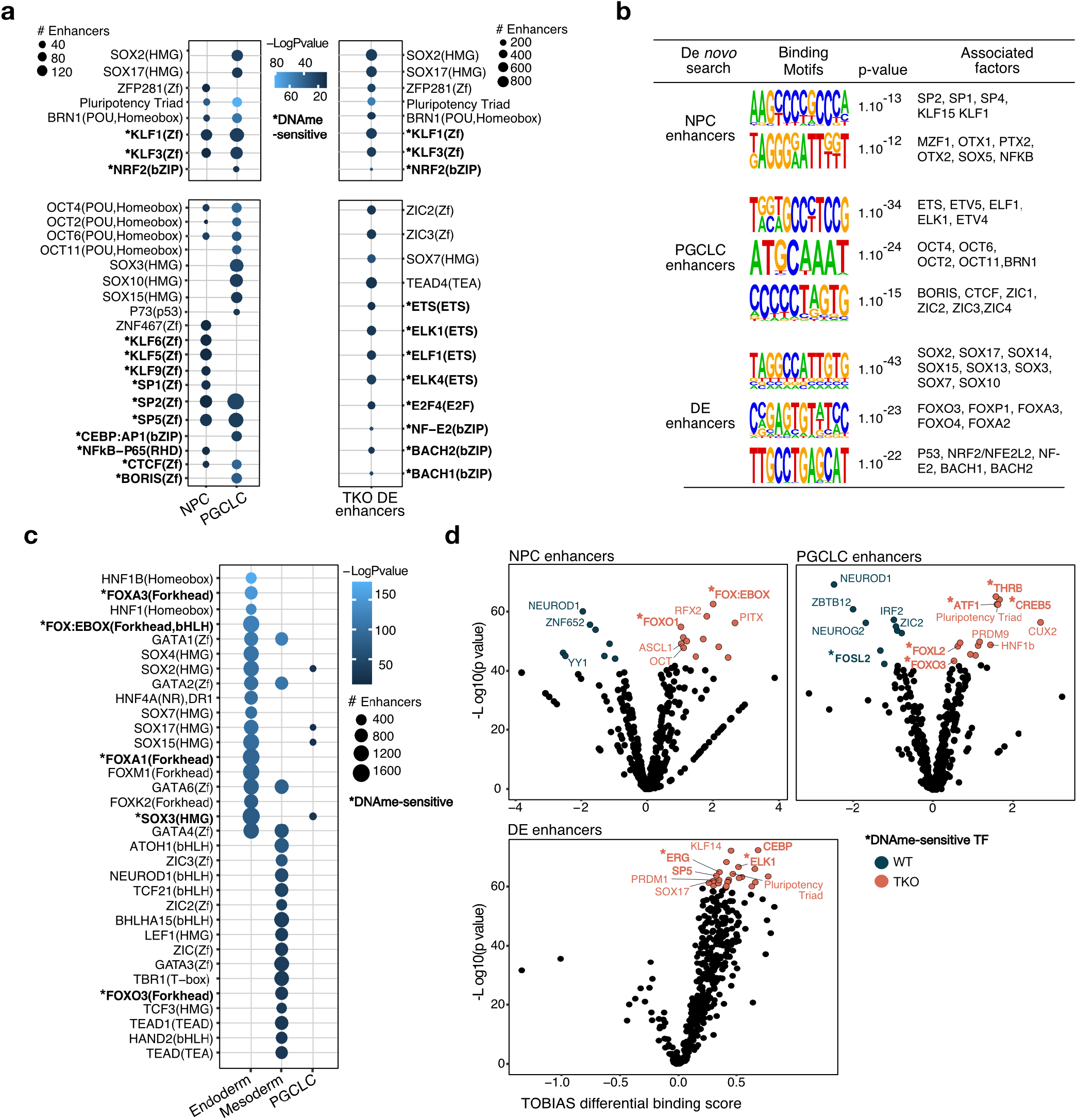
Differential mobilization of DNA methylation-sensitive TFs at NPC and PGCLC enhancers. **a,** Bubble plot of HOMER analysis for top 20 TF binding motifs at NPC (*n*= 168), PGCLC (*n*= 350) and all DE (*n*= 1,339) putative enhancers. Factors highlighted in bold with an (**\***): DNA methylation-sensitive TFs or regulated by DNA methylation. Color gradient represents -log(pvalue), circle size shows the number of enhancers bearing the binding motifs. **b,** HOMER detection of motifs at NPC, PGCLC and all DE enhancers. *De novo* motifs and known associated TFs are represented, with their associated p-value. **c,** same as in a), but with endoderm and mesoderm enhancers. **d,** TOBIAS footprint analysis of predicted differential binding score of TF occupancy at NPC, PGCLC and all DE enhancers in D4 TKO over WT EpiLCs.

To prove that these TFs are actually mobilized on relevant enhancers in TKO EpiLCs, we performed Transcription factor Occupancy prediction By Investigation of ATAC Signal (TOBIAS) ^49^. TOBIAS detects drops in accessibility at TF binding motifs as a sign of physical occupancy of the corresponding TFs (described as footprints). Looking at NPC, PGCLC and DE enhancers in D4 TKO EpiLCs, we observed an increased binding score for motifs associated with DNA methylation-sensitive factors (FOXO, SP, CEBP…), or lineage regulators such as ASCL1 or the Pluripotency Triad (OCT4, SOX2 and NANOG) (Fig. 6d). These TOBIAS-identified TFs belonged to gene clusters that dynamically gained expression during priming (clusters 1, 2 and 4) in both WT and TKO EpiLCs (Extended Data Fig. 8c and Supplementary Table 6).

As a whole, this indicates that DNA methylation-sensitive and/or lineage regulators bind NPC and PGCLC enhancers more strongly or more frequently among cells during TKO EpiLC differentiation, while these enhancers normally slowly gain DNA methylation during WT EpiLC differentiation. Importantly, we found these characteristics to be specific to this enhancer category, and were not observed at meso/endoderm enhancers. Moreover, our defined NPC and PGCLC enhancers demonstrated similar chromatin and DNA methylation dynamics, as well as TF regulation, suggesting a previously unappreciated coordination of the neural and germline fate during priming.

## DISCUSSION

By systematically comparing the ability of DNA methylation-free ESCs to respond to various differentiation routes, we provide here novel insights into the role of DNA methylation on key developmental bifurcations. DNA methylation appeared unnecessary for the priming program, as well as for the induction of neural progenitors during somatic differentiation. DNA methylation was also not required for forming primordial germ cells, but rather seemed to exert a temporal control over germline opportunity. Whether DNA methylation also restrains the window of germline competence *in vivo* is an important question, which we were not able to address due to the developmental delay of chimeric embryos containing TKO cells and the difficulty to stage match with WT chimerae.

It was recently reported that treating embryoid bodies with demethylating agents promoted the adoption of PGC identity by delaying the exit from naive pluripotency ^31^. However, in genetically impaired DNA methylation-free TKO cells, we did not find evidence that the extended time-window for germline specification was due to improper extinction of naive pluripotency genes or to delayed priming. Interestingly, extended temporality for PGCLC specification was previously observed when EpiLCs were exposed to alpha-ketoglutarate (aKG), a mitochondrial oxidative phosphorylation (OxPhos) metabolite, which can promote DNA hypomethylation in naive stem cells ^50,51^. However, we did not observe misregulation of OxPhos-associated genes in TKO EpiLCs (data not shown), indicating that DNA methylation does not regulate germline specification through metabolism.

Instead, we propose that DNA methylation may close the temporal window of primordial germ cell specification by progressively decommissioning key enhancers during naive-to-primed transitioning (Figure 7). Indeed, we observed that putative enhancers of PGCLC genes maintained abnormally high chromatin accessibility and H3K27ac enrichment during TKO EpiLC differentiation. Similarly, NPC-related enhancers also remained more strongly engaged in TKO EpiLCs, in association with preferred neural differentiation. The chromatin relationship between PGCLC and NPC enhancers is intriguing, but we could link it to similar sequence composition, and in particular, binding motifs for common DNA methylation-sensitive and/or developmental TFs that are present during EpiLC differentiation and fail to decommission from these enhancers in absence of DNA methylation. It is noteworthy that some of these TFs, such as OCT4, CREB and ATF, are known to favor histone acetylation through direct association with CBP/p300 ^52–54^.

**Fig. 7.**
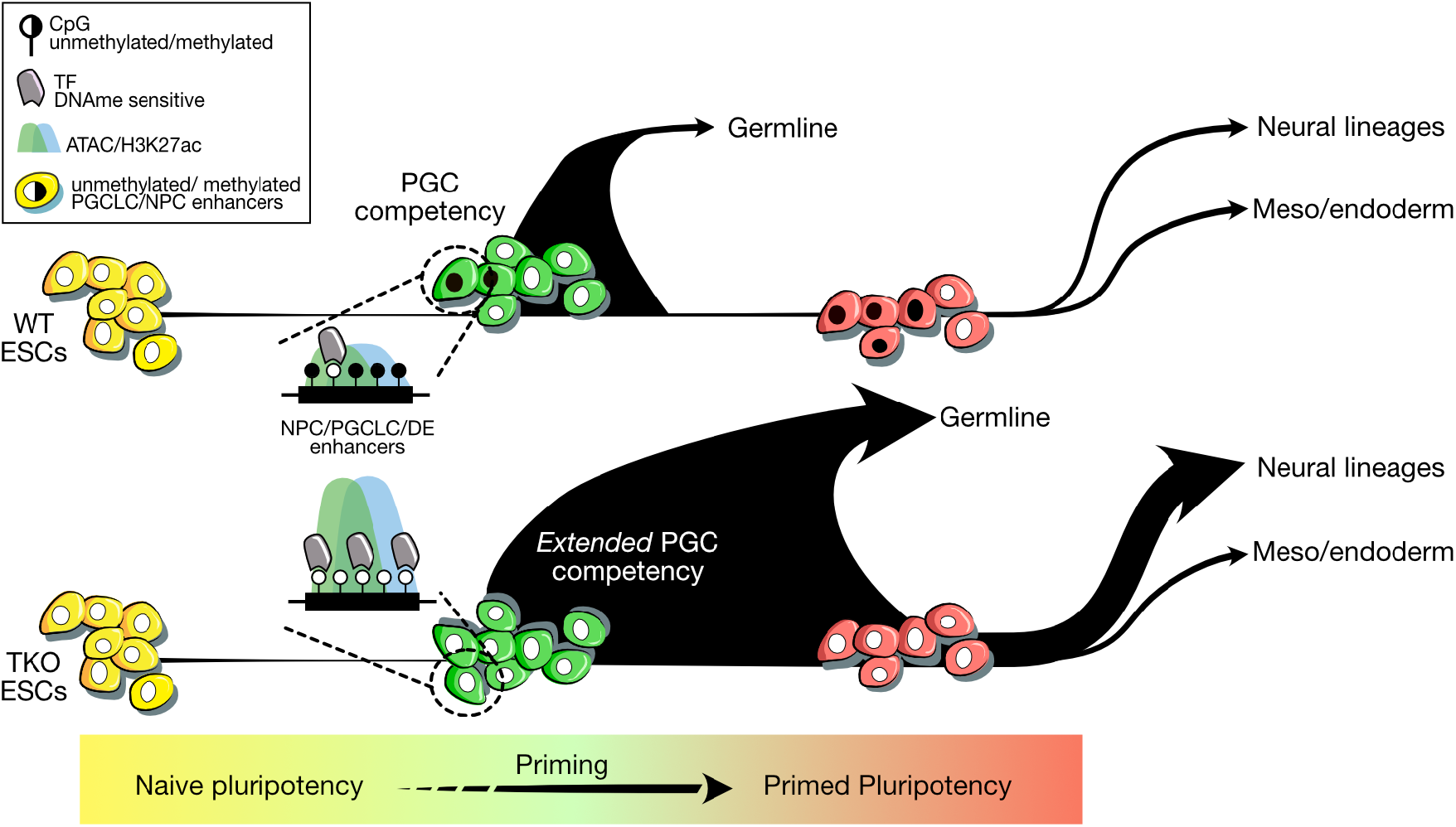
Graphical summary for DNA methylation role in tempering neural and germline fates as default differentiation routes. In naive hypomethylated ESCs, chromatin is globally relaxed, including at NPC and PGCLC enhancers. During priming, WT EpiLCs heterogeneously gain DNA methylation at a subset of NPC and PGCLC putative enhancers. This is associated with restricted chromatin accessibility, H3K27ac loss, and decreased binding of DNA methylation-sensitive TFs, hence balancing cell fate diversity by limiting neural and germline fates. In contrast, in absence of DNA methylation, TKO EpiLCs fail to temper their neural and germline “hyper competent” state. This is associated with retention of permissive chromatin signatures and DNA methylation-sensitive TF binding at NPC and PGCLC enhancers, thus skewing cell fate decisions towards these lineages during priming.

Several evidence suggest that cell fate choices could be epigenetically primed in the enhancer landscape of progenitors cells ^55,56^. In naive and primed ESCs, somatic lineage-associated enhancers harbor heterogenous levels of DNA methylation and around 3% of them were estimated to be regulated by methyl-sensitive TFs ^20^. Variegating DNA methylation reflects the competing activity of DNA methylation writers and erasers—namely DNMTs and ten-eleven translocation (TET) enzymes—during priming ^15,17,57^, which seems to be particularly pronounced at enhancers of the ectoderm lineage during epiblast development ^45^ Accordingly, recent single cell analyses reported that embryos lacking DNMT1 have over-represented neuro-ectoderm-associated cell types ^37,58^. Our dynamic and time-resolved culture system therefore further emphasizes the role for DNA methylation in tempering neural fate, while revealing for the first time that germline fate is also subject to DNA methylation-based restriction during priming.

Why would the neural and germline lineages be co-regulated? It is interesting to note that the proteome and transcriptome of brain and testis tissues are strikingly similar in adult mice and humans ^59^. On an evolutionary perspective, such brain-testis axis could reflect the central role played by these two tissues in speciation events ^60^. This similarity does not only reflect common tissue organization between these two organs, as transcriptomic convergence is also prominent when comparing isolated neurons and sperm cells versus other cell types ^59,61^. What our work implies is that coordinated regulation of germ cells and neural cells could have an early embryonic origin—at the time of their specification—through a common set of DNA-methylation sensitive TFs present in the epiblast. Further experiments should be carried out to challenge this intriguing, early functional link and understand the importance of such common regulation of the neural and germline lineages by DNA methylation.

## Acknowledgements

We are grateful to all Bourc’his lab members for their continuous support. We thank J. Hackett for the piggyback vector containing the H2B::tdTomato reporter, M. Cohen-Tannoudji for critical reading of the manuscript, M. Borenzstein for advices in generating PGCLCs and I. Kucinski for the SMARTseq2 RNA-seq design. We acknowledge the Core Cytometry platform and the ICGex NGS platform of Institut Curie (supported by grants ANR-10-EQPX-03, Equipex and ANR-10-INBS-09-08, France Génomique) and the Cell and Tissue Imaging Platform-PICT-IBiSA (member of France-Bioimaging, ANR-10-INBS-04) of the Genetics and Developmental Biology Dpt (UMR3215/U934) of Institut Curie. The laboratory of D.B. is part of the LABEX DEEP (ANR-11-LABX-0044, ANR-10-IDEX-0001-02). This work was supported by the Fondation Bettencourt Schueller and the Fondation pour la Recherche Médicale (FRM Team Label). M.S. was supported by PhD fellowships from Ministère de l’Enseignement Supérieur et de la Recherche and from Fondation pour la Recherche Médicale.

## Author contributions

M.S., M.V.C.G. and D.B conceived the study. Most experiments were performed by M.S. A.T. performed the bioinformatic analyses on ATAC and CUT&RUN, P.G. on scRNA-seq, and M.S. on bulk RNA-seq data. E.D.M. performed NPC experiments. M.A. performed library generation. J.I. and F.M. performed chimeras and whole-mount embryo IF. S.K. and B.G performed scRNA-seq. M.W. generated the WGBS data. D.B. and M.S. interpreted the data and wrote the manuscript. All authors reviewed and approved the final manuscript.

## Data availability

The data supporting the findings of this study are available in the main text and the supplementary materials. All sequencing data will be made publicly available after publication of this article in a peer-reviewed scientific journal.

## MATERIAL AND METHODS

### Mouse embryonic stem cell lines

Mouse male embryonic stem cells (ESCs, E14 TG2a background derived from the 1291*0la* strain) were used to generate a triple knockout (tKO) for the three DNA methyltransferase genes *Dnmt1*, *Dnmt3A* and *Dnmt3B*. The strategy for CRIPSR-Cas9 editing of the catalytic PCN/Q loop of the DNMT MTase domain was previously described in Domcke et al., 2015.

### ESC culture

ESCs were cultured in feeder-free conditions at 37°C in 5% CO_2_ atmosphere. Mycoplasma-free status was assessed using the MycoplasmaCheck service from Eurofins.

#### Serum-LIF culture medium

ESCs were cultured on 0.1% gelatin-coated plates with serum-LIF medium (Glasgow medium (Sigma), 2mM L-glutamine (Gibco), 0.1mM MEM non-essential amino acids (Gibco), 1mM sodium pyruvate (Gibco), 15% Fetal Bovine Serum (FBS), 0.1mM β-Mercaptoethanol and 1000 U/ml leukemia inhibitory factor (LIF, Chemicon)). Cells were counted on a Beckman Cell Counter Vi-Cell XR and passed with TrypLE (Trypsin replacement enzyme, Gibco) every 2-3 days.

#### 2i + Vitamin C culture medium

To reduce DNA methylation levels, ESCs were grown for at least two weeks on fibronectin-coated plates (1/60 dilution in PBS) in 2i+VitC medium (N2B27 (50% Neurobasal medium and 50% DMEM/F12 (Gibco) complemented with 2mM L-glutamine, 0.1mM β-Mercaptoethanol, N2 supplement (Gibco) and B27 serum-free supplement (Gibco)), complemented with 1000U/ml LIF, 2i (3μM Gsk3 inhibitor and 1μM of MEK inhibitor), and vitamin C 100µg/ml final (Sigma)) and passed with accutase (GIBCO) every 2-3 days.

#### Epiblast-like cell (EpiLC) differentiation

To initiate EpiLC differentiation, 2i+VitC- grown ESCs were first split with accutase (Gibco) and plated on fibronectin-coated 6-well/plates at a density of 0.20×10^6^ cells/well. The next day, the medium was replaced with AF medium: N2B27 medium supplemented with 12ng/ml Fgf2 (R&D) and 20ng/ml Activin A (R&D), 1% Knock-out Serum Medium (Gibco), to drive EpiLC differentiation. Cells were harvested at Day 2, 4, and 7 of differentiation, with an additional splitting step at D4 with accutase to reduce cell density.

#### Neuronal progenitor cell (NPC) differentiation

For monolayer NPC differentiation, serum-LIF-grown ESCs were split with accutase onto laminin-coated 6-well/plates (Sigma, 10μg/ml in PBS) at a density of 0.75×10^4^ cells/well in plain N2B27 medium. Cells were cultured up to 10 days with daily medium change.

#### Embryoid body differentiation

Embryoid bodies were generated by dissociating 2i+VitC-grown ESCs with TrypLE and 3×10^6^ cells were seeded in uncoated B10 plates in serum without LIF to allow the formation of cell aggregates. Medium was changed every 2 days.

#### Meso-endoderm differentiation

Meso-endoderm differentiation was adapted from ^23^. Briefly, serum-LIF-grown ESCs were split with accutase onto fibronectin-coated plates (Sigma, 10μg/ml in PBS) at a density of 1×10^4^ cells/ well in 6-well/plates in N2B27 basal medium supplemented with 10 ng/ml Activin A and 3 µM of GSK3 inhibitor CHIR99021. Cells were cultured for 6 days, and the medium was changed every two days.

#### PGC-like cell differentiation

After 40h, 48h ou 96h of EpiLC differentiation, cells were resuspended in PGCLC basic medium: GMEM (GIBCO), KSR 15% (GIBCO), Non-Essential Amino Acid (GIBCO), Sodium Pyruvate 1mM (GIBCO), L-Glutamine 2mM (GIBCO), Penicillin-Streptomycin 100U/ml (GIBCO), β-Mercaptoethanol 0.5mM (GIBCO). Then EpiLCs were distributed into ultra-low adhesive U-bottom 96-well plates at a density of 2.5×10^3^ cells/well in 100μl of PGCLC medium complemented with cytokines for PGCLC induction (PGCLC basic medium, 0.5 µg/ml BMP4 (R&D system), 0.5 µg/ml BMP8 (R&D system), 0.1 µg/ml stem cell factor (SCF, R&D system), 50 ng/ml epidermal growth factor (EGF, R&D system), 1000 u/ml mouse LIF). After 4 days of culture, embryoid bodies were harvested and dissociated with 10mM EDTA in PBS+3%FBS for subsequent FACS analyses.

### Cell transfection and cell line isolation

All vectors were generated using Gibson cloning from ^62^. For sgRNA cloning, pX330 (Addgene #42230) or pX459 (Addgene #62988) plasmids were digested with *BbsI* restriction enzyme (NEB) and annealed with double-stranded sgRNA oligos using QuickLigase (NEB). The H2B::tdTomato-containing piggyBac vector was a kind gift from Dr. Jamie Hackett (EMBL Rome).

Transgenic ESC lines were generated by nucleofection using Amaxa 4D nucleofector (Lonza) using 3-5× 10^6^ cells with 1-3µg of vectors and sgRNA/Cas9 plasmids. Cells were then seeded in serial dilutions in 0.1% gelatin-coated B10 plates in serum-LIF. After two days of recovery, cells underwent puromycin (1μg/ml, Life Technologies) or hygromycin B (200µg/ml Sigma) selection for 2 or 4 days, respectively. After selection, the culture medium was changed for serum-LIF only and individual colonies were picked and screened by PCR genotyping.

### Live-imaging for apoptosis analysis

To quantify apoptosis, cells were grown on a 96-well plate and placed in the incubated chamber of an Incucyte ZOOM Microscope (Sartorius). EpiLC differentiation medium was added and supplemented with 5µM of Caspase 3/7 Green dye (Sartorius) to detect for apoptosis in live cells. Medium was changed every day, brightfield and green fluorescence images were taken every 2 hours for 96 hours. Counting of dead cells was made using IncuCyte ZOOM Software.

### Fluorescence activated cell cytometry and sorting

Cells were collected, dissociated with TrypLE, resuspended with PBS/FBS 3% supplemented with DAPI (1µg/ml) to discriminate dead cells. For PGCLC specification quantification, 2.5µL/10^5^ PGCLCs were incubated with Anti-CD61-PE (Biolegend #104307) and Anti-SSEA1-eFluor660 (eBioscience #50-8813-42) antibodies in PBS/FBS 3% for 30 min on ice in the dark, before being analyzed on the ZE5 Flow cytometer (Biorad). PGCLC sorting was performed on a Sony SH800. List of all antibodies references and working dilutions are available in Supplementary Table 8.

### Karyotyping

ESCs were treated with KaryoMAX Colcemid (Gibco) in PBS for 2h at 37°C. After dissociation using TrypLE, cells were resuspended in 3ml of hypotonic solution (1 volume of FBS+3 volumes of H2O) for 10min at 37°C. After centrifugation, cells were fixed with 3ml of Carnoy solution (Acetic acid diluted ¼ in 100% ethanol) and incubated at room temperature (RT) for 30 min. Cells were then dropped from ∼50cm height onto glass slides. Slides were air-dried and DNA was stained with DAPI (1µg/ml). Chromosomes were imaged with an upright widefield microscope (Leica) and around 100 cells were analyzed per cell line on Fiji for counting.

### Western blot analysis

Proteins were extracted by sonicating cell pellets after treatment with protease inhibitors (Aprotinin, Leupeptin, Pepstatin and PMSF) and BC250 buffer containing 1M TRIS pH8, 20% glycerol, and 0.5M EDTA. After centrifugation (10 min-9000g at 4°C), the supernatant was denatured at 95°C for 10min and equal quantities of proteins were transferred on a 4%-12% Bis-Tris Gel (NuPage). After electrophoresis, the gel content was transferred on 0.45µm nitro-cellulose membranes (GE Healthcare). After blocking with 5% milk and PBS/NP40 (PBS+0.3% NP40), membranes were incubated overnight with antibodies diluted 1:5000 in 1% milk in PBS/NP40. After washes in PBS/NP40, membranes were incubated 2h at RT with HRP-conjugated secondary antibodies. Revealing was performed by 5 min incubation with LumiLight^Plus^ (Roche) substrate on a Chemidoc MP Imaging system (BioRad). List of antibody references and working dilutions are available in Supplementary Table 8.

### LUMA analysis

Global DNA methylation levels were assessed using the Luminometric Methylation Assay (LUMA) ^63^ from WT and TKO ESCs in serum-LIF medium, and WT ESCs cultured in 2i medium complemented with VitC. Briefly, 500ng of genomic DNA was digested with *MspI*+*EcoRI* and *HpaII*+*EcoRI* (NEB) in parallel reactions at 37°C for 4h, followed by heat inactivation at 65°C. *EcoRI* is included as an internal reference. Filling of the genome-wide protruding ends of the restriction digestions were quantified in a pyrosequencing reaction (PyroMark Q48 autoprep, Qiagen, dispensation order: ACTCGA) and all results were analyzed with the associated software (Q48 Autoprep Software).

### Liquid chromatography-mass spectrometry (LC-MS)

LC-MS-based quantification of methyl-cytosines was performed on 1 μg of DNA degraded to nucleosides with nuclease P1 (Roche), snake venom phosphodiesterase (Worthington) and alkaline phosphatase (Fermentas). An equal volume of isotopic standard mixture (15N3-C (Silantes), 2H3-5mC (TRC) and self-synthesized 15N3-5hmC, 15N3-5fC and 15N3-5caC were added to the nucleoside mixture and injected for LC-MS/MS analysis as described previously Schomacher et al., 2016. Quantitative analysis was performed on an Agilent 1290 Infinity Binary LC system (Agilent technologies) using a ReproSil 100 C18 column (Jasco) coupled to an Agilent 6490 triple quadrupole mass spectrometer (Agilent technologies).

### Dual Luciferase assay

Dual luciferase assays were performed using Promega DLR kit (#E1910) according to the manufacturer’s instructions. WT and TKO ESCs were cultured in 2i+VitC for transfection at D0, or cultured in AF medium for 2 days for EpiLC transfection. Cells were co-transfected with pGL3EV/pCAGRLuc (Empty control vector) and pGL3DE/pCAGRLuc (distal enhancer) or pGL3PE/pCAGRLuc (proximal enhancer). Briefly, 1.6μg of each plasmid was diluted in a mix of 500μl of OptiMEM media (GIBCO) and 4μl of Lipofectamine 2000 (Thermoscientific) per well. Culture medium was replaced with the transfection mix for half a day, and then completed with 500μl of AF medium. The next day, medium was changed for AF or 2i+VitC and cells were allowed to grow one more day before harvest and lysis using PLB buffer for luminescence analysis on CalrioStar (BMG Labtech).

### ESC Reversion assay

After 2, 4 and 7 days of EpiLC differentiation, cells were split in accutase, and plated in fibronectin-coated 6-well plates at the density 1×10^5^ cells/well in 2i+VitC medium. Colonies from D2, D4 or D7 EpiLCs were grown for 4, 7 and 10 days, respectively, before being stained for alkaline phosphatase activity. Briefly, cells were rinsed with PBS and fixed with ethanol 100% for 10 min at RT. After 3 PBS rinses, cells were incubated at 37°C for 10 min with FastBCIP/NBT (Sigma #B5655) dissolved in MiliQ water, and air-dried. Pictures were taken and analyzed on Fiji to count AP-stained reverted ESC colonies.

### Chimerae

Mice were hosted on a 12h light/12h dark cycle at 22 +/- 2°C ambient temperature and 40-70% humidity, with free access to food and water, in the pathogen-free Animal Care Facility of the Institut Curie (agreement C75-05-18). All animal experiments were performed following European (Directive 2010/63/EU) legislation and were approved by the ethics committee of the Institut Curie CEEA-IC #118 and by French Ministry of Research (APAFiS#30920-2021040622279416-v1).

#### Chimera aggregation

Eight-week-old B6D2F1 (C57BL/6J × DBA2) females, were superovulated by intraperitoneal (i.p.) administration of 5 IU of Pregnant Mare Serum Gonadotropin (PMS013 Centravet) followed by an additional i.p. injection of 5 IU Human Chorion Gonadotropin (CH0003 Centravet) 48 hours later. Females were mated to a stud male of the same genetic background. Morula embryos were collected and washed in M2 medium (Sigma M7167) and cultured in Cleave medium (Cook K-RVCL) at 37 °C under 5% CO2. Meanwhile, low passage WT and TKO ESCs containing the H2B::tdTomato reporter were thawed and cultured in 2i medium. Cells were dissociated by accutase in small clumps of 5-10 cells. Right before aggregation, the zona pellucida of morulae was removed by Tyrode’s acid treatment (Sigma T1788-100ML), before being washed with M2 medium. Each morula-stage embryo was placed into a depression well performed using an aggregation needle (BLS DN-09) on a plastic plate. A dissociated ESC clump was added close to the embryo to allow aggregation and development to the blastocyst stage. Integrity and proper integration of transgenic ESCs in the inner cell mass were assessed with a fluorescence inverted microscope (Zeiss Axio Vert A.1) before re-implantation into pseudo-pregnant foster mothers (NMRI). Embryos were harvested at E7.5 and E8.5 for whole-mount immunostaining.

### Immunofluorescence

For whole-mount embryo immunostaining, E7.5 and E8.5 embryos were fixed in 4% PFA for 30 min at RT. Then, embryos were washed in PBT (PBS+0,1% Triton) and permeabilized for one hour at room temperature in PBS+0,5% triton. Blocking was performed by adding 0.2% BSA and 5% Donkey serum in PBT for 30 minutes. Primary antibodies diluted in blocking solution were incubated for 72 hours at 4°C with the embryos, then three washes of 15min at RT were performed. Similarly, secondary antibodies were diluted in blocking solution and incubated for 3 hours at RT. After the final washes, embryo mounting was made through increasing percentages of glycerol as follow: 2.5% (5 minutes), 5% (5 minutes), 10% (10 minutes), 20% (15 minutes), 50% (15 minutes), and DTG (Glycerol, DABCO 2.5%, 50mM Tris pH8.6) (15 minutes). Images were obtained using a Zeiss LSM800 inverted confocal microscope. List of antibody references and working dilutions are available in Supplementary Table 8.

### RT-qPCR

RNA extraction was performed on cell pellets using standard Trizol (Life Technologies)-chloroform protocol. RNA pellet was treated with DNase before clean up with RNeasy Mini kit (QIAGEN). To generate cDNA first-strand, a minimum of a 100ng of RNA was reverse transcribed using random priming with SuperScript III (Life Technologies). The design of the qPCR primers was made using Primer3 program). List of all primers used are available in Supplementary Table 7. Amplification reactions were conducted on the Viia7 thermal cycling system (Applied Biosystems) using SYBR Green Reagent (Thermo Fisher Scientific). Expression levels were normalized to housekeeping genes using ΔΔCT method.

### Bulk RNA sequencing

For NPC and EpiLC bulk RNA sequencing, total RNA was isolated as described above. RNA-seq libraries were prepared from a minimum of 200 ng of DNase-treated total RNAs with the TRuSeq Stranded mRNA protocol (Illumina). Between 59M and 130M reads were sequenced per sample in a 100bp paired-end format using a NovaSeq 6000 (Illumina) platform. For PGCLC bulk RNA sequencing, libraries were generated from 50ng of isolated of DNase-treated total RNA using SMARTer Stranded Total RNA-seq Kit for mammalian Pico Input. Between 38M and 45M reads were sequenced per sample.

### Single-cell RNA sequencing

Single cells were index-sorted individually by FACS (SONY SH800) into wells of a 96-well PCR plate containing 2.3 µl of lysis buffer (200U/µL SUPERase-IN (LifeTechnologies) and 10% Triton X100 in RNase-free water). A scRNA-seq SMARTseq2 protocol was performed as previously described^65^. The Illumina Nextera XT DNA kit was used to prepare libraries that were sequenced on the Illumina HiSeq 4000 (paired-end 50bp). Samples from all cell lines were included in each sequencing lane, to control for technical lane effects. Only cells with over 7500 detected reads assigned to genes were kept in the analysis.

### CUT&RUN

The CUT&RUN protocol was optimized from ^66^. Concanavalin A beads (Polysciences) were first activated in 1ml of Binding Buffer (20mM HEPES-KOH, 10mM KCl, 1mM CaCl_2_, 10mM MnCl_2_). Then, between 200 000 and 300 000 cells were aliquoted in 1ml of Wash Buffer (20mM HEPES-KOH, 150mM NaCl, 0.5mM Spermidine (Sigma) and 1X Complete EDTA-free (Roche) protease inhibitor cocktail) and 20µL of activated beads were added to the cells, followed by 5 min incubation at RT on a rotating wheel. Cells were collected on a magnetic rack; the supernatant was replaced by 400µL of primary antibody diluted to 1:200 in Wash Buffer complemented with 2mM EDTA and 0.02% Digitonin (Millipore). Cells were incubated for one hour at RT. Samples were then washed twice in Dig-Wash Buffer (Wash Buffer + 0.02% Digitonin) for 15 min at RT and incubated with 1:400 home-made pA-MNase in Dig-Wash for 15 min at RT. List of all antibodies references and working dilutions are available in Supplementary Table 8.

After two more washes, cells were resuspended in 150µl of Dig-Wash and cooled-down for five min at 0°C. For targeted digestion, CaCl_2_ was added to the cells to a final concentration of 2mM followed by 30 min incubation on ice. The reaction was stopped by adding 150µl of 2X STOP Buffer (340mM NaCl, 20mM EDTA, 4mM EGTA, 0.02% Digitonin, 100ng/ml RNase1, 250ng/ml glycogen) and incubating the samples at 37°C for 20 min before centrifugation at 4°C 16000g for 5 min. Samples were placed on a magnetic rack and supernatant was recovered in a low binding eppendorf tube. Following addition of 0.1% SDS and 0.17mg/ml Proteinase K, samples were incubated at 70°C for 30 min. Purified DNA was obtained by phenol/chloroform extraction and precipitated with 100% Ethanol incubation at −20°C for at least 20 min. The DNA pellet was washed in 85% ethanol, spun down and air-dried before being resuspended in 25µl of 1mM Tris-HCl pH8.0.

Library preparation was made according to the manufacturer’s instructions (Accel NGS 2S DNA library kit, Swift Biosciences). For library amplification, 12 PCR cycles were applied as followed: 98°C for 10 sec, 60°C for 15 sec and 68°C for one minute. Library size and quantification was assessed on Agilent 4200 Tapestation. CUT&RUN libraries were sequenced on NovaSeq (Illumina) using PE 100bp run, with biological duplicates for all samples. Between 25M and 75M reads were sequenced per sample.

### ATAC sequencing

The ATAC protocol was optimized from ^67^. First, 1×10^7^ ESCs were mixed with 2.5×10^6^ *Drosophila* S2 cells, and resuspended and incubated in 1ml Lysis Buffer (50mM KCl, 10mM MgSO_4_.7H_2_0, 5mM HEPES, 0.05% NP40) for 5 min at RT. After a spin at 4°C 1500g for 5 min, the supernatant was discarded and nuclei pellet was resuspended in ice-cold RS Buffer (10mM NaCl, 10mM Tris-HCl, 3mM MgCl_2_). Nuclei were counted using Beckman Coulter ViCell XR cyto-counter, to ensure nuclei integrity. Tagmentation was performed on 50 000 nuclei per sample, with TDE1 Tn5 transposase (Nextera), at 37°C for 30 min. The reaction was stopped by the addition of PB Buffer and tagmented DNA was eluted using minElute columns from Reaction clean-up Qiagen Kit. We used 50ng of tagmented genomic DNA isolated from the same nuclei preparation to control for Tn5 transposition bias. Libraries were generated by PCR amplification (maximum of 12 cycles) as previously described in Buenrostro et al., 2013 using custom-made barcoded primers. List of all barcodes used are available in Supplementary Table 7. Libraries were cleaned on minElute columns and quality was assessed using Agilent 4200 Tapestation. Sequencing was performed on NovaSeq platform (Illumina) using PE 100bp run. Between 40M and 91Mreads were sequenced per sample.

### Definition of PGCLC-NPC-EpiLC-specific genes

To define PGCLC and NPC specific genes, differential gene expression was compared between NPCs at D8, PGCLCs after 40h of EpiLCs induction, and primed EpiLCs at D7. Genes were declared as NPC-specific or PGCLC-specific if FDR < 0.05 and log2(CPM) > 1 when compared to the other two states (comparisons were NPC vs PGCLC/EpiLCD7, and PGCLC vs NPC/EpiLCD7). To define EpiLC-specific genes, the same strategy and thresholds were applied, using EpiLC at D2 instead of EpiLC at D7 to capture transiently expressed “formative” epiblast genes (comparisons were EpiLC D2 vs PGCLC/NPC). This list of defined genes can be found in Supplementary table 2.

### WGBS

Whole Genome Bisulfite sequencing was produced as described previously^69^. Briefly, genomic DNA from EpiLC at D3 was isolated using the GenElute Mammalian Genomic DNA Miniprep Kit (Sigma) with RNase treatment. Whole-Genome Bisulfite Sequencing libraries were prepared from 50ng of bisulfite-converted genomic DNA using the EpiGnome/Truseq DNA Methylation Kit (Illumina) following the manufacturer instructions. Sequencing was performed in 100 pb paired-end reads at a 30X coverage using the Illumina HiSeq2000 platform.

### WGBS data analysis

Whole-genome bisulfite sequencing was analyzed as described previously ^69^. Briefly, reads generated in this study or recovered from available data-sets were treated as follow. The first eight base pairs of the reads were trimmed using FASTX-Toolkit v0.0.13 (hannonlab.cshl.edu/fastx_toolkit/index.html). Adapter sequences were removed with Cutadapt v1.3 (code.google.com/p/cutadapt/) and reads shorter than 16 bp were discarded. Cleaned sequences were aligned onto the Mouse reference genome (mm10) using Bismark v0.12.5 ^70^ with Bowtie2-2.1.0 ^71^ and default parameters. Only reads mapping uniquely on the genome were conserved. Sequencing statistics can be found in Supplementary Table 1. Methylation calls were extracted after duplicate removal. Only CG dinucleotides covered by a minimum of 10 reads were conserved for the rest of the analysis. We completed this D3 EpiLC WGBS dataset with D0 (available at GSE71593) and D7 (available at GSE121405) time-points datasets produced previously in our lab and available publicly.

### Bulk RNA-seq analysis

Paired-end read trimming was made using trim Galore (v065) or NPC and PGCLC data sets, and Atropos (github.com/jdidion/Atropos) for EpiLC data sets. Next, alignment was made using Kallisto pseudo-aligner (v0.46.2). Gene annotation was obtained from Ensembl (EnsDb.Mmusculus.v79). Sequencing statistics can be found in Supplementary Table 1.After EdgeR (v3.30.3) CPM normalization, differential gene expression analysis was made using voom transformation from Limma R package (v3.44.3). P-Values were determined using Limma and adjusted with the BH correction, cutoff was set up at 0.05. Threshold for differentially expressed genes were set at the FDR<5% and |log2FC| >1.

### Single-cell RNA-seq analysis

Paired-end reads were trimmed with trim Galore (v0.6.5) in order to remove adapter sequences and Ns nucleotides from both side of the read. Cleaned reads were aligned onto a concatenated genome (the Mouse reference genome mm10 + fasta file with the ERCC spike-in sequences) with STAR (v2.7.5a) reporting random alignment with at most 6 % of mismatches. ERCCs, genes and TE families-based quantification was performed with featureCounts (v1.5.1) using a concatenated file with repeatMasker, ERCC spike-in and Gencode vM25 annotations. Filtering of cells was performed with the following criteria: cells with less than 7500 genes with at least one read and cells with more than 2% of mitochondrial reads or more than 10% of ribosomal reads were filtered out. On the remaining cells, the raw counts were normalized with the SCTransform from Seurat (v4.0.1). PCA was performed on the 10000 most variable genes defined by the vst method. UMAP was performed on the 10 first principal components with 10 neighbors, a minimal distance of 0.1 and a spread of 1. Cells clusters were defined with the FindNeighbors and FindClusters functions from Seurat on 5 first principal components with 14 neighbors and a resolution of 0.4. The list of specific markers for the single-cell clusters is available in Supplementary table 3. Pseudotime was inferred with the R package slingshot (v1.8.0) on the 10 first principal components.

### CUT&RUN analysis

Paired-end reads were trimmed with trim Galore (v0.4.4) in order to remove adapter sequences and Ns nucleotides from both side of the read. Cleaned reads were aligned onto the Mouse reference genome mm10 with bowtie2 (v2.2.9) in end-to-end and very sensitive mode. Duplicated reads were removed using Picard (v2.6.0). Sequencing statistics can be found in Supplementary Table 1. Bigwig files for UCSC genome browser and heatmaps were created with deepTools (v2.5.3). Sliding windows approach was used to define enriched regions with csaw R package (v1.24.3). Window size of 150bp and 5 kb was used for H3K27ac and H3K27me3 samples with a spacing interval of 38bp and 1250bp respectively. Window-level counts were quantified using the *windowCounts* function, discarding blacklisted regions (ENCODE, accession ENCFF547MET), with mapping quality threshold of 20. For each genotype (WT and TKO), non-insteresting windows were filtered out using a global background enrichment (*filterWindowsGlobal* function) with large bins (750bp for H3K27ac and 25kb for H3K27me3 samples). Normalization factors were calculated using the large bins and used for normalization of enriched windows to remove composition biases. Differential binding analysis was performed with csaw. Windows that are less than 100bp and 1kb (for H3K27ac and H3K27me3 samples respectively) apart were merged into regions. Regions with a FDR<5 % were declared as differentially bound. List of H3K27ac and H3K27me3 CUT&RUN regions is in Supplementary Table 4.

### ATAC-seq analysis

Paired-end reads were trimmed with trim Galore (v0.4.4) in order to remove adapter sequences and Ns nucleotides from both side of the read. Sequencing statistics can be found in Supplementary Table 1. Cleaned reads were aligned onto the Mouse reference genome mm10 with bowtie2 (v2.2.9) in end-to-end and very sensitive mode. Mitochondrial reads were removed from the rest of the analysis. Duplicated reads were removed using Picard (v2.6.0). Bigwig files for UCSC genome browser and heatmaps were created with deepTools (v2.5.3). Enriched regions were defined using MACS2 (v2.1.1) with a q-value threshold of 0.01, discarding the ones that overlap with blacklisted regions (ENCODE, accession ENCFF547MET). Overlapping regions between biological replicates were used for the following analyses. Region-level counts were quantified using the *windowCounts* function with mapping quality threshold of 20. Normalization factors were calculated using 10kb-long bins to remove composition biases. Differential binding analysis was performed with csaw. Regions that are less than 500bp apart were merged. Regions with a FDR<5 % were declared as differentially bound. List of ATAC regions can be found in Supplementary Table 5.

### Integration of CUT&RUN and ATAC-seq data

Genes annotated in close proximity (less than 5kb to the TSS) to differentially bound regions were used for gene ontology enrichment analysis with the R clusterProfiler package (*enrichGO* function). Overlaps between differentially bound regions defined in different time points and epigenetics marks were performed using intervene (v0.6.4). Enhancers were defined as the co-occurrence of ATAC and H3K27ac-enriched regions that are distal from promoter regions (+/- 2kb around the TSS). PGCLC and NPC enhancers were defined as enhancers that are +/- 50kb from the TSS of PGC and NPC genes that were defined as previously explained. The list of putative PGCLC and NPC enhancers can be found in Supplementary Table 6. Quantification of NPC, PGCLC, endoderm, mesoderm enhancers and bivalent genes was performed with featureCounts (v1.5.1). Bivalent genes in EpiLCs and mesoderm/endoderm enhancers were originally defined in ^37,72^. Enrichment of known motifs and *de novo* motif finding were performed using HOMER (v4.11) ^43^ with the default database. Transcription factor activity, also called footprinting, was assessed using the software TOBIAS (v.0.13.3) ^49^. Tn5 bias correction was first performed using the ATACorrect module. Then footprinting scores were calculated using the corrected cut sites with ScoreBigwig module. Differential binding score between conditions was estimated with BINDetect module with the HOMER motif database.

### Softwares

Statistical analyses and graphs were performed with two-tailed unpaired t-test using GraphPad Prism (v9.4). Microscopy image analyses were made possible using Fiji (ImageJ2 v2.3.0). Transgene constructions were made possible with Geneious (v7.1). Figures were made with Affinity Designer (v1.8.6).

## EXTENDED DATA FIGURES

**Extended data Fig. 1.**
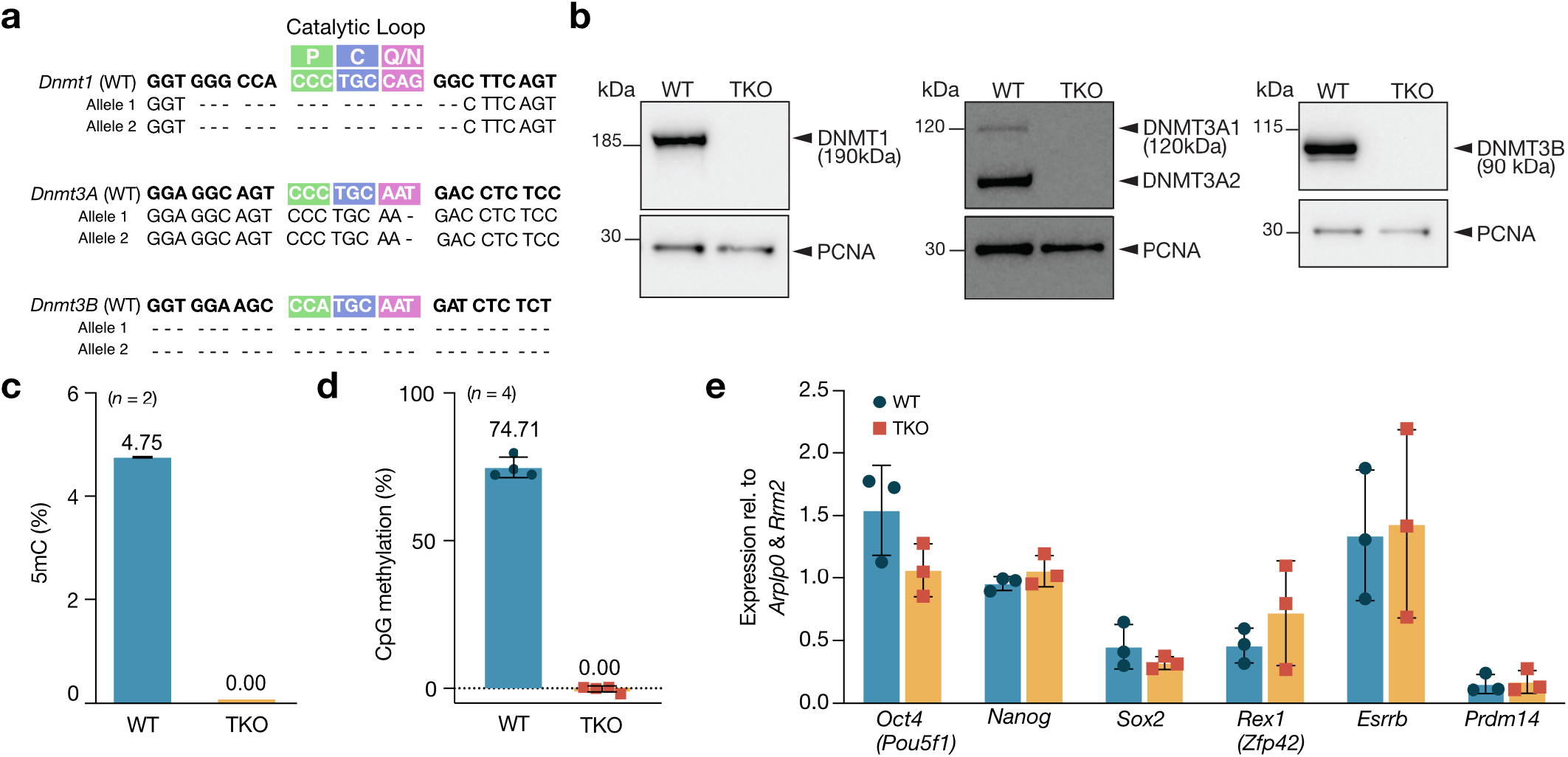
Characterization of DNA-methylation-free *Dnmt*-triple KO (TKO) ESCs. **a,** Sequencing results for the E14 TKO ESC line showing allelic homozygous deletions of *Dnmt* genes after catalytic motif targeting by CRISPR-Cas9. **b,** Immunoblotting showing DNMT1, DNMT3A and DNMT3B in WT and TKO ESCs grown in serum/LIF. PCNA is used as a loading control. **c,** Barplot showing LC/MS quantification of methylated cytosines in serum/LIF-grown WT (blue) and TKO (orange) ESCs. Data shown are mean ± SD in technical duplicates. **d,** LUMA assay of genome-wide CpG methylation of serum/LIF-grown WT and TKO ESCs. Data shown are *n=* 4 biological replicates with mean ± SD. **e,** Expression of pluripotency genes measured by RT-qPCR in 2i/LIF-grown WT and TKO ESCs. Data shown are mean ± SD from biological triplicates, ΔCT values are normalized to *Arplp0* and *Rrm2*.

**Extended data Fig. 2.**
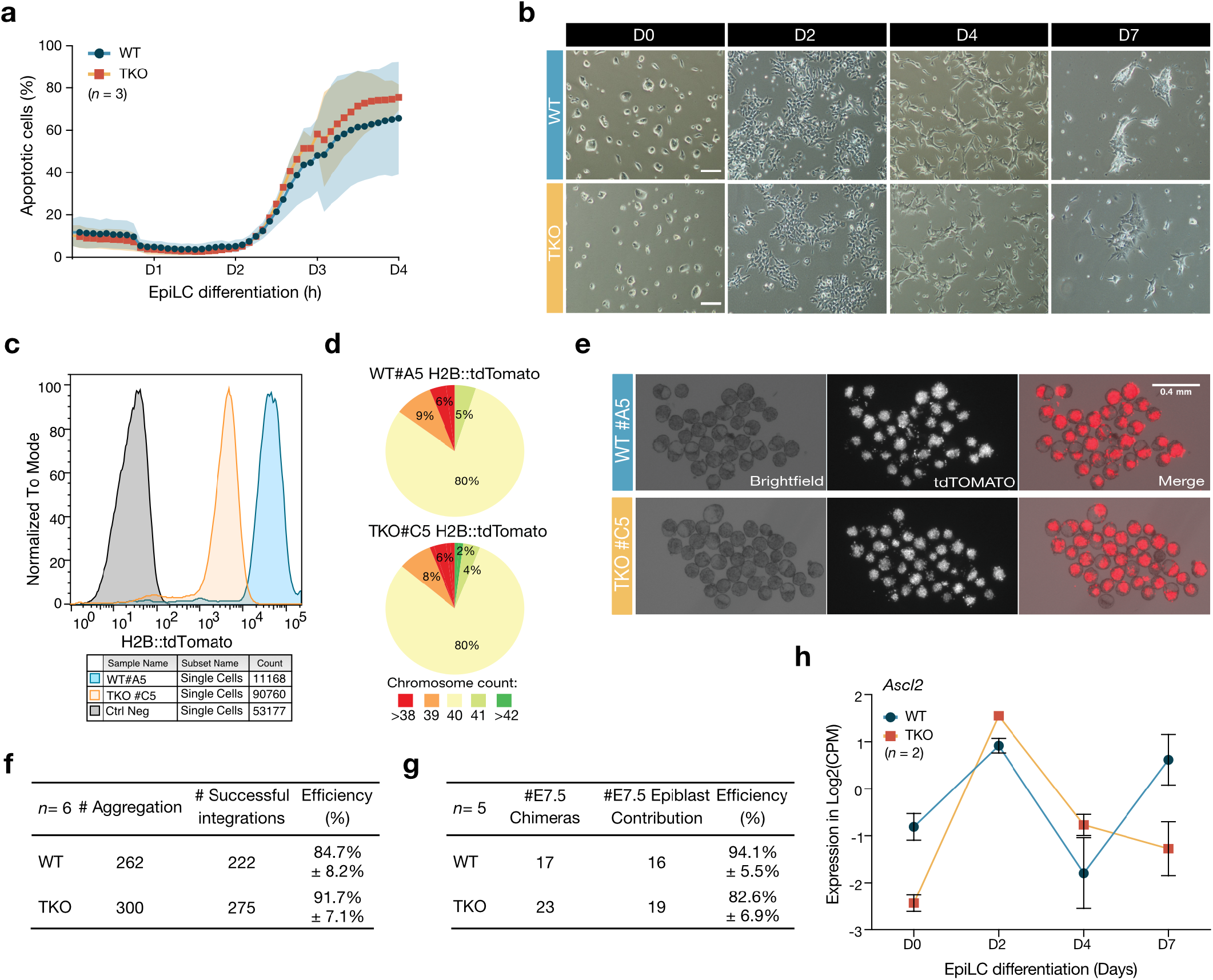
TKO EpiLCs acquire functional priming features *in vitro* and *in vivo*. **a,** Apoptosis rate of WT and TKO cells, measured by live staining of Caspase 3/7 on Incucyte. Data shown are from biological triplicates with mean ± SD. **b,** Representative images of WT (blue) and TKO (orange) cells during EpiLC differentiation. Scale: 100µm. **c,** Flow cytometry analysis of H2B::tdTOMATO reporter expression in WT clone #A5 (blue) and TKO clone #C5 (orange), compared to untransfected negative control cells. **d,** Karyotyping and chromosome counting in WT #A5 and TKO #C5 reporter cell lines, that were selected for chimera aggregation. **e,** Representative microscopy pictures after 24h of culture of aggregates of host-morula/ H2B::tdTOMATO ESCs. The merge reveals contribution of both WT and TKO ESCs to the blastocyst ICM. Scale: 400µm. **f, g,** Statistics for aggregation experiments. **(f)** Six aggregation experiments (*n=* 6) were performed, totalizing 202 aggregates of WT #A5 and 231 aggregates of TKO #C5 reporter ESCs with host morulae. Integration efficiency, measured by the number of blastocysts with correctly integrated reporter ESCs after 24h of culture. (**g**) Five chimera assays (*n=* 5) were recovered after re-implantation of the aggregates in pseudo-pregnant females. Despite low yield and some non-chimeric embryos (94.1% and 82% of chimeras from WT and TKO aggregates, respectively), all recovered chimerae showed contribution of reporter cell lines in the embryonic epiblast only. **(h)** Expression of trophectoderm marker *Ascl2* during EpiLC differentiation. Expression is shown as normalized Log2 CPM counts (average between biological duplicates).

**Extended data Fig. 3.**
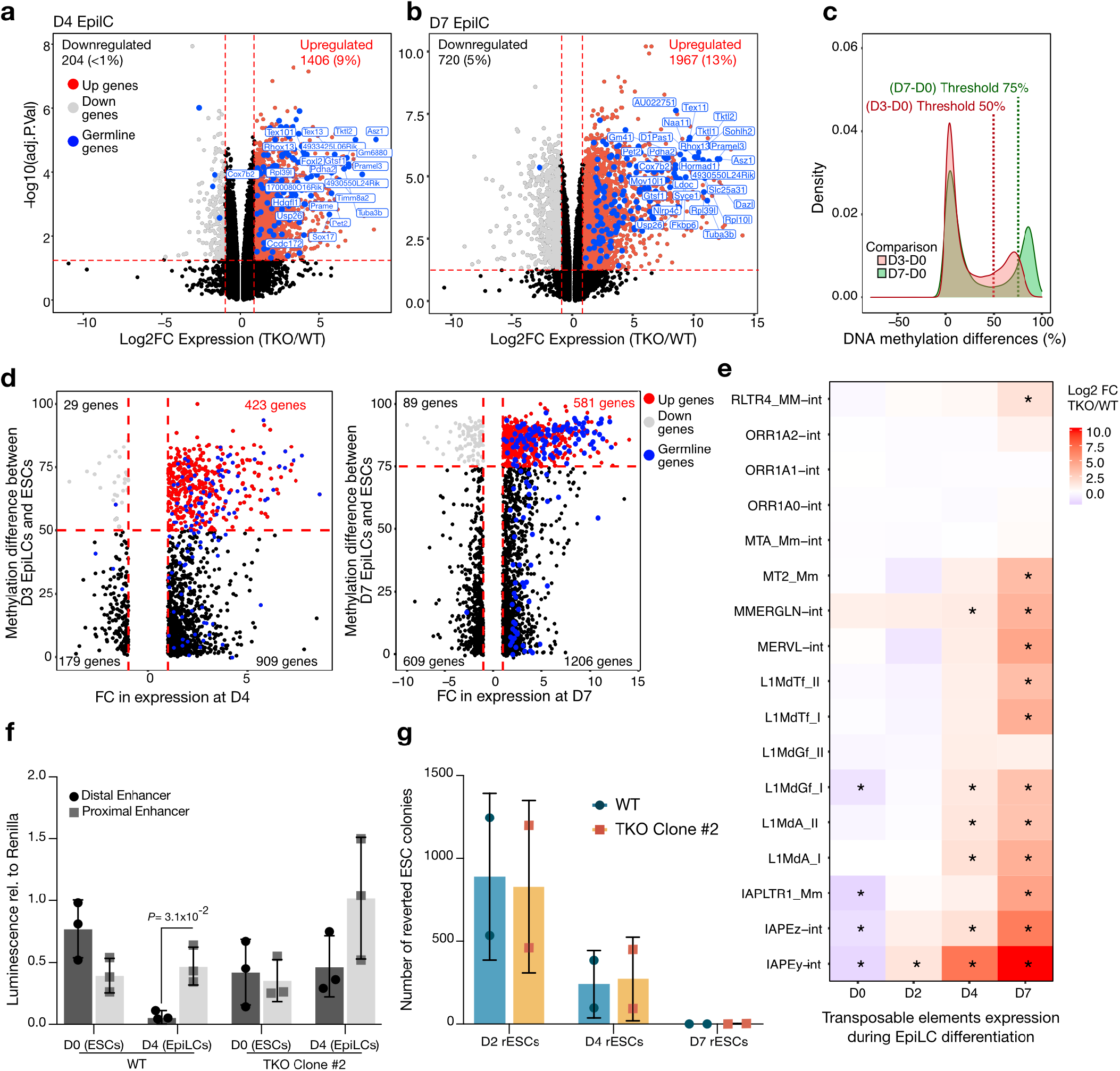
TKO EpiLCs upregulate germline genes and some classes of transposable elements. **a, b,** Volcano plots showing differential expression in TKO over WT EpiLCs at D4 (**a**) and D7 (**b**) in Log2FC versus -log10(adj. pvalues). Red: up-regulated genes; Grey: down-regulated genes; Blue: germline genes. Thresholds were set up at FDR <5% and Log2(FC)>1. Data obtained from biological duplicates. **c.** Density plot showing differences in DNA methylation between D0, D3 and D7 of EpiLC differentiation at WT cells promoters (<10kb from TSS) as measured by WGBS ^34^. Threshold for regions defined as differentially methylated (DMRs) was arbitrarily set-up at 50% between D0-D3 and 75% between D0-D7. **d**, Scatter plots showing the correlation between DNA methylation gain at gene promoters during WT EpiLC differentiation and expression in Log2(FC) in TKO versus WT EpiLCs. Left: DNA methylation level changes at D3 over D0 ESCs and expression changes at D4. Right: DNA methylation level changes at D7, expression changes at D7. Red: up-regulated genes with a DMR promoter; Blue: germline genes. **e,** Heatmaps of transposable element expression showing normalized fold-change in TKO over WT cells during EpiLC differentiation. Black stars (*) represent significance (FDR<1%, Log2FC>1). Expression was obtained for each condition from the average between biological duplicates. **f,** Barplot representing Luciferase activity from the distal (dark grey) and proximal (light grey) *Oct4* enhancers in WT and in an independent TKO clone #2 in D0 ESCs and D4 EpiLCs. Data are mean ± SD from biological triplicates (two-tailed unpaired student t-test). **g,** Barplot displaying numbers of reverted ESC colonies upon 2i+VitC medium replacement at D2, D4 and D7 of EpiLC differentiation in WT and in an independent TKO clone #2. Data are from biological duplicates with mean ± SD.

**Extended data Fig. 4.**
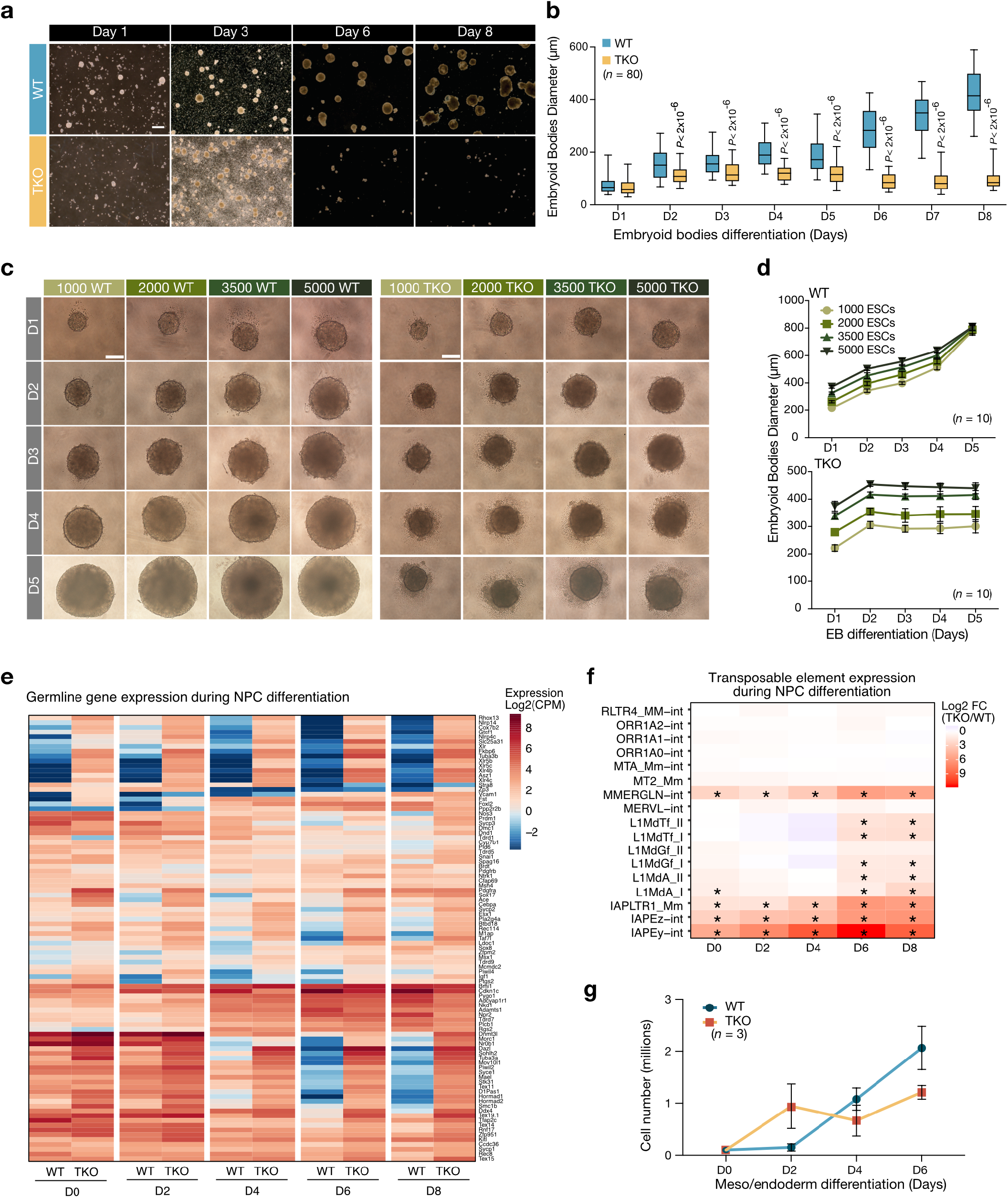
TKO Cells can under directed, but not undirected, somatic differentiation. **a**, Brightfield images of embryoid bodies (EBs) generated from WT (blue) and TKO (orange) ESCs cultured in serum for 8 days. Scale: 500µm. **b**, Boxplot representation of EB diameter across differentiation. Boxes represent 5-95 confidence interval, *n=* 80 EBs counted/genotype/timepoint (two-tailed unpaired student t-test). **c**, Brightfield images of EBs derived from WT and TKO ESCs at different seeding densities (1000, 2000, 3500 and 5000 cells). Scale: 200µm. **d**, EB diameter size curve showing the effect of increasing initial cell density during a five-day long differentiation. Data shown are biological replicates (n=10) with mean ± SD. **e**, Heatmaps of germline markers showing normalized Log2 CPM counts during NPC differentiation. Expression is obtained for each condition from the average between biological duplicates. **f,** Heatmap of transposable elements showing Log2(FC) in TKO over WT cells during NPC differentiation. Black stars (*) represent significance (FDR<1%, Log2FC>1). **g**, Growth curve of WT and TKO ESCs subject to meso/endoderm differentiation. Data are mean ± SD in biological triplicates.

**Extended data Fig. 5.**
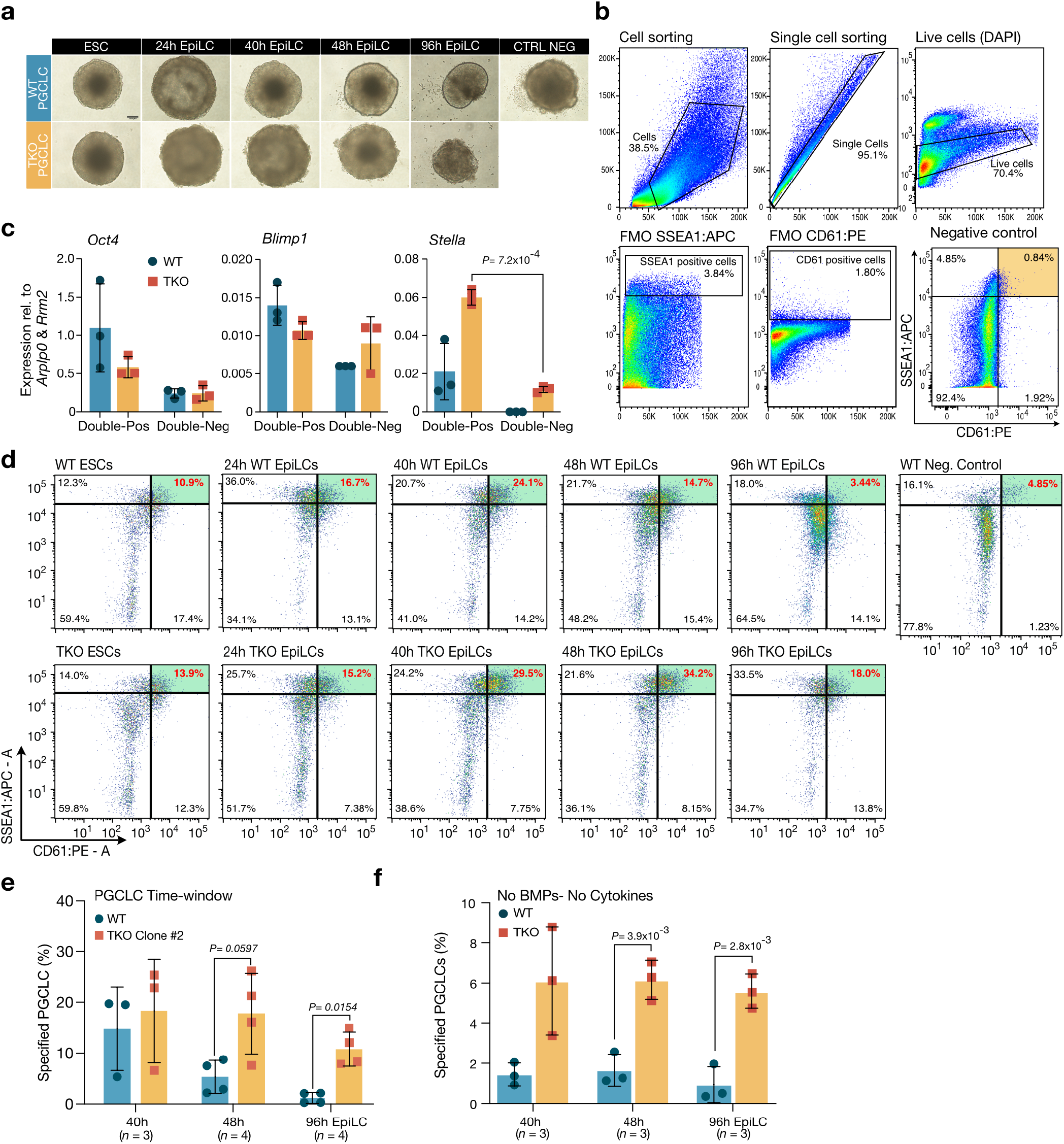
TKO EpiLCs can undergo germline differentiation *in vitro*. **a,** Representative brightfield images of cultured PGCLC aggregates at D4 generated from WT and TKO cells cultured in presence or absence of PGCLC-inducing cytokines. Scale: 250µm. **b,** FACS strategy: after dissociation from aggregates grown under PGCLC-inducing conditions, single cells were selected based on SSC/FSC. Then, single cells were identified by comparison between SSC-area/SSC-height. Next, live cells were selected based on DAPI integration. Finally, PGCLCs were identified for positive staining with SSEA1 and CD61 surface markers. Thresholds were set by fluorescence minus one (FMO) conditions and negative control, i.e. cells cultured without PGCLC-inducing cytokines. **c,** Expression of PGCLC markers measured by RT-qPCR in WT (blue) and TKO (orange) PGCLCs. Data shown are mean ± SD from ΔCT values from biological triplicates, normalized to *Arplp0* and *Rrm2* (two-tailed unpaired student t-test). **d,** Representative FACS plot of SSEA1-Pos and CD61-Pos WT and TKO PGCLCs generated from ESCs and EpiLCs at different time points. Cell percentages are indicated in each quarter. **e,** Barplot showing the percentage of specified WT and TKO PGCLCs generated from EpiLCs at different times, in an independent E14 TKO clone #2. Data shown are mean ± SD from biological replicates (*n*=3 or 4, two-tailed unpaired student t-test). **f,** Barplot showing percentage of specified WT and TKO PGCLCs generated from ESCs and EpiLCs at different times, in absence of pro-germline cytokines. Data shown are mean ± SD from biological replicates (*n*=3, two-tailed unpaired student t-test).

**Extended data Fig. 6.**
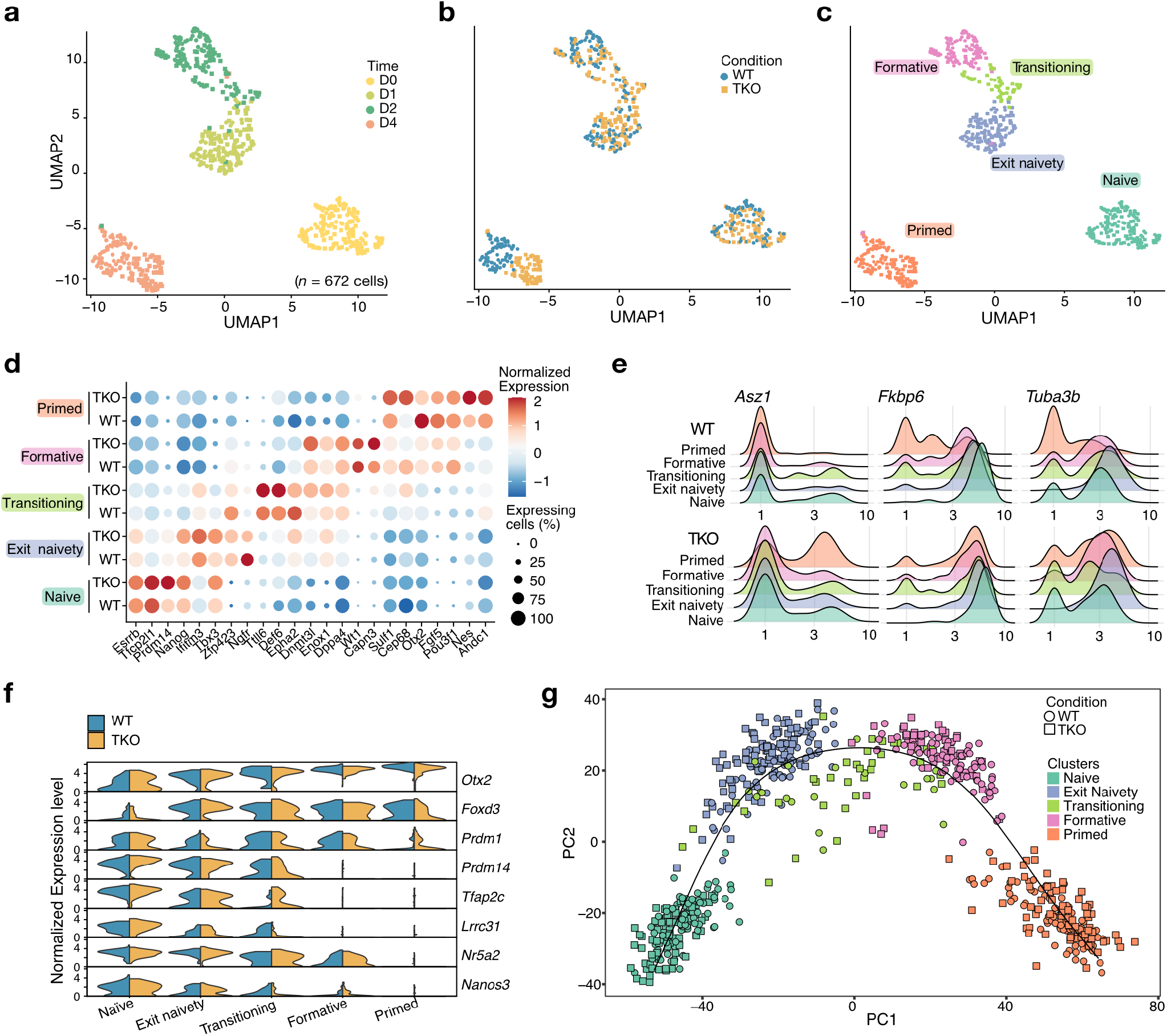
Increased ability of TKO EpiLCs for germline specification does not originate from a “germline-competent” subpopulation. **a, b, c,** UMAP dimensionality reduction of scRNA-seq data from WT and TKO EpiLCs highlighting (**a**) Timing of differentiation, (**b**) Condition (WT in blue, TKO in orange) and (**c**) unbiased cell clustering for a total *n*= 672 cells, from two biological duplicates. **d,** Dotplot of expression levels of key pluripotency and priming markers in each cluster detected for WT and TKO cells. Diameter of the dots represents the percentage of cell expressing the marker, colors represent average expression levels. **e,** Ridgeplot displaying germline gene expression (in Log scale) in each identified cluster for WT and TKO cells. **f,** Violin plot showing expression of PGC fate regulators in WT and TKO EpiLCs in each of the identified clusters. **g,** PCA dimensionality reduction for pseudotime trajectory of WT and TKO EpiLCs highlighted for identified clusters during EpiLC differentiation.

**Extended data Fig. 7.**
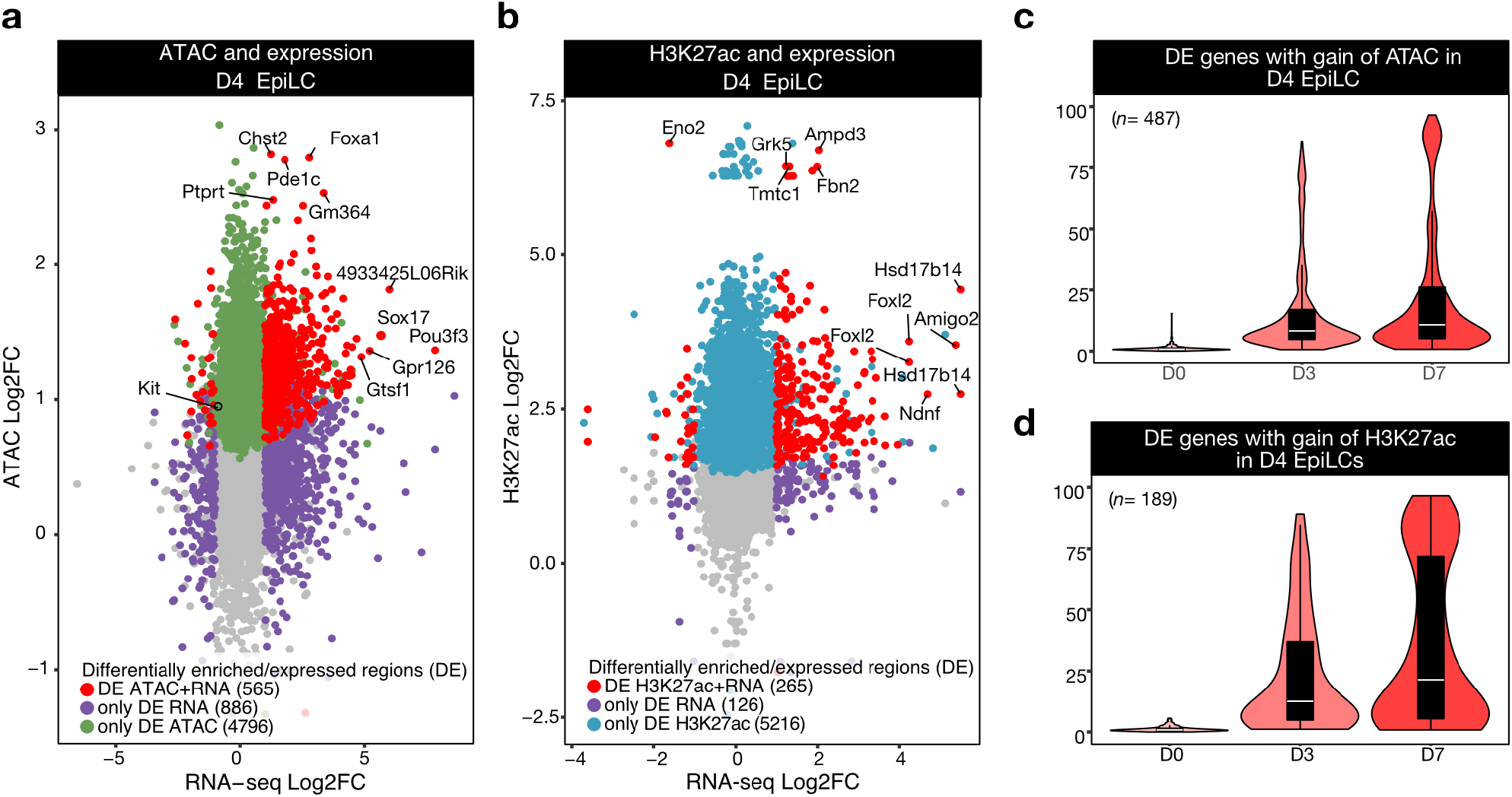
Uncoupling between chromatin changes and expression changes in TKO EpiLCs. **a, b,** Scatterplots showing Differential Enrichment in log2 Fold-change in (**a**) ATAC and (**b**) H3K27ac vs Differential Expression in log2 Fold-change in RNA-seq in D4 EpiLCs. Regions were annotated to genes at close proximity (<5kb from TSS). Colored dots depict the status of expression and/or enrichment. Numbers between brackets represent the number of regions marked by (**a**) ATAC in green and (**b**) H3K27ac in blue in D4 EpiLCs. For differentially expressed (DE) genes with differentially enriched (DE) regions in close proximity (red dots), the top 10 DE genes are displayed. **c, d,** Violin plots showing the dynamics of methylated CpGs in during WT EpiLC differentiation (D0, D3, D7), around promoter (<1kB) of DE genes associated with differential enrichment in (**c**) ATAC and (**d**) H3K27ac in D4 TKO EpiLCs, using previously published WGBS datasets ^34^. Data shown are the median with upper and lower hinges corresponding to 75 and 25% quantile.

**Extended data Fig.8.**
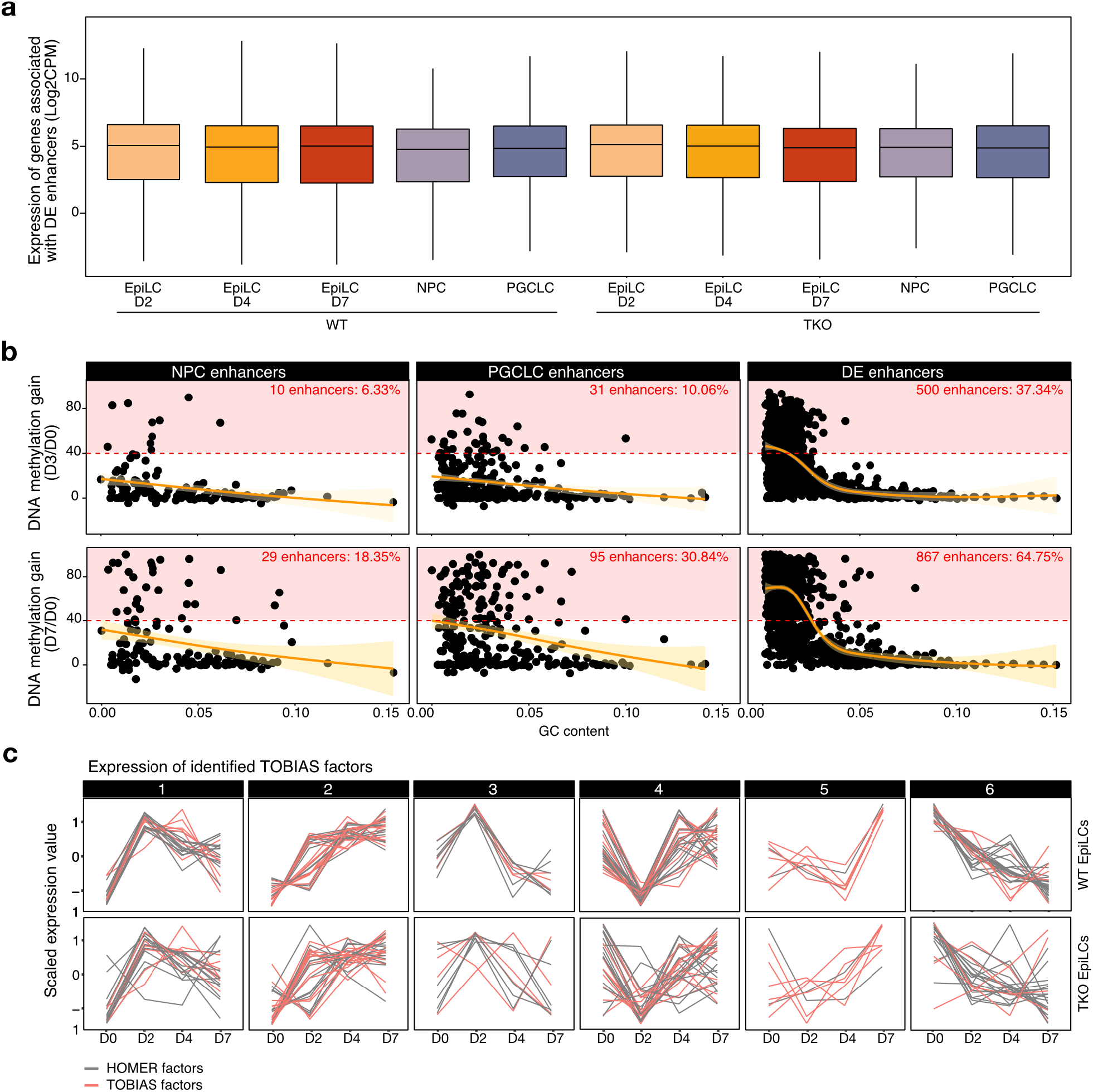
Chromatin changes at NPC, PGCLC and DE enhancers in TKO EpiLCs. **a,** Expression of genes in the vicinity of all identified DE putative enhancers (within 50kb) in WT and TKO EpiLCs, NPCs and PGCLCs (40h). **b,** Scatterplot showing DNA methylation gain between D0/D3 (Top) or D0/D7 (Bottom) in WT EpiLCs at NPC, PGCLC and DE enhancers, according to GC content. Thresholds have been defined (red dotted lines) for subsets of enhancers gaining high DNA methylation levels during EpiLC differentiation. Orange curves correspond to generalized additive mode smoothing method with standard error (yellow). **c,** Clusters showing the transcriptional dynamics of TFs (scaled expression values) identified via HOMER (grey) and TOBIAS (pink) during WT and TKO EpiLC differentiation.

